# Novel mouse model of cerebral microbleeds created by Crispr/Cas9-mediated *Col4a1* deletion in adult brain microvessels

**DOI:** 10.1101/2025.06.22.660900

**Authors:** Hyunmi Kim, Yeojin Seo, Heekyung Kho, Shivani Shailendrakumar Singh, Jina Lee, Hohyeon Lee, Ji-Won Hwang, Tae-Ryong Riew, Seungyon Koh, Jun Young Choi, Hyun Woong Roh, Sang Joon Son, Gi Tae Kim, Sung Kweon Cho, Hyun-Seok Jin, Seon-Yong Jeong, Kang In Lee, Jae Young Lee, Byung Gon Kim

## Abstract

Cerebral small vessel disease is a leading cause of cognitive decline and stroke in the elderly, with cerebral microbleeds (CMBs) as one of the key imaging biomarkers. Our understanding of its pathophysiology remains limited due to the lack of appropriate animal models. We report a novel mouse CMB model created by disrupting collagen IV, a core component of the vascular basement membrane (BM), specifically within brain microvessels. Targeted deletion of *Col4a1* was achieved in adult mice using brain endothelial-specific AAV vectors with CRISPR/Cas9. MRI revealed numerous CMBs with distributions similar to those of human CMBs. CMB burden increased progressively over six months following *Col4a1* deletion in a dose-dependent manner, accompanied by cognitive decline and motor incoordination. Histological examination revealed hemosiderin deposits corresponding to MRI-detected CMBs without evidence of macroscopic hemorrhage or white matter lesions, while ultrastructural analysis demonstrated significant BM thinning in *Col4a1*-depleted microvessels. Analysis of human MRI and genomic data identified significant associations between CMB susceptibility and genetic variants in *TIMP2*, an endogenous inhibitor of the matrix-degrading enzyme MMP2, underscoring the clinical relevance of our model. These findings establish a direct causal relationship between microvessel COL4A1 and CMB, suggesting that dysregulated collagen IV homeostasis in BM underlies CMB development.

## Introduction

Cerebral small vessel disease (CSVD) is a frequent cause of progressive cognitive decline in the elderly, often accompanied by other neurological dysfunctions such as gait disturbance, extrapyramidal signs, and sphincter dysfunctions ^1,2^. Recent advancements in magnetic resonance imaging (MRI) technology have facilitated the detection of brain pathologies linked to changes in small cerebral blood vessels, which include lacunar infarctions, white matter hyperintensities, widened perivascular spaces, and cerebral microbleeds (CMBs) ^3^. Following stroke, patients with CSVD often show deteriorated functional recovery, a higher risk of future vascular events, and reduced post-stroke survival ^4,5^. For example, in patients with ischemic stroke or transient ischemic attack, the presence of CMBs increases the risk of recurrent stroke, particularly intracerebral hemorrhage (ICH) ^6^. Moreover, CSVD frequently coexists with various neurodegenerative diseases, especially Alzheimer’s disease, and can worsen cognitive dysfunctions and other neurological disabilities associated with neurodegeneration ^2,3^. Despite these significant clinical burdens, our understanding of the underlying pathogenesis of this disease entity is very limited, in part due to a lack of adequate animal models ^7^.

The basement membrane (BM) in the cerebral small blood vessels is located at the interface between the circulatory system and the central nervous system (CNS). It plays a vital role in the maintenance of the blood-brain barrier (BBB) and the structural integrity of the blood vessels ^8^. The BM is made up of a three-dimensional network composed of four major glycoproteins, with collagen IV isoform serving as a stabilizing polymer network ^8,9^. The collagen IV in the brain vascular BM has been linked to various vascular pathologies. In humans, mutations in the *COL4A1* gene can cause perinatal cerebral hemorrhage and porencephaly in infants ^10,11^. Adults with *COL4A1* mutations not only present with ICH but also exhibit CSVD pathological phenotypes, including white matter hyperintensities, widened perivascular spaces, lacunar stroke, and CMBs ^11–14^. Genetic analysis of mice with *Col4a1* mutants revealed that the mutation causes vascular fragility and increases the likelihood of ICH in both newborn and adult mice ^10,11^. However, the potential contribution of Col4a1 in the brain microvessels to various CSVD phenotypes in sporadic adult patients remains to be further investigated.

The purpose of this study was to explore whether a genetic perturbation of the *Col4a1* specifically in the brain microvessels could produce a range of CSVD phenotypes in adult mice. To deplete the *Col4a1* gene in adult brain microvessels, we administered CRISPR/Cas9 associated single guide RNA (sgRNA) targeting *Col4a1* using engineered adeno-associated virus (AAV) vector that were shown to specifically target brain endothelial cells (BECs) ^15^ in transgenic mice that overexpress Cas9. Our findings showed that CRSPR/Cas9-mediated deletion of *Col41* in the adult brain microvessels led to the development of numerous CMBs in a highly penetrable manner, without any evidence of other CSVD phenotypes or macro-ICH. This striking CMB phenotype suggests a direct link between changes in collagen IV and the development of sporadic CMBs. Furthermore, analysis of genetic data from human subjects with available magnetic resonance imaging (MRI) revealed associations between susceptibility to CMB and variants of genes involved in maintaining collagen IV, highlighting the clinical relevance of our CMB model.

## Results

### CRISPR/Cas9-mediated editing of *Col4a1* gene in mouse BECs

To generate sgRNA targeting the mouse *Col4a1* gene, nine sgRNAs targeting coding regions were designed and cloned into a Cas9-expressing vector. Plasmids containing Cas9 and the candidate gRNA sequences targeting the mouse *Col4a1* gene were transfected into mouse fibroblast cell line (NIH-3T3), which express *Col4a1*. To evaluate the insertion or deletion (indel) mutations mediated by Cas9 at the targeted sites, targeted deep sequencing was performed. We found that sgRNAs #1, #4, and #5 targeting exon 1 of mouse *Col4a1* exhibited higher indel efficiency compared to other gRNAs (Fig. S1A). We performed secondary screening with sgRNAs (#1, #4, and #5) and found that sgRNA #4 exhibited the highest out-of-frame indel efficiency, while sgRNA #5 resulted in slightly greater reduction in the mRNA expression of *Col4a1* (Fig. S1B, C). To select the best performing sgRNA, we used primary brain endothelial cells (BECs) obtained from Cas9-expressing transgenic mice. When both sgRNAs were transfected into the BECs using lentivirus, we found that sgRNA #4 was more effective than sgRNA #5 in reducing the amount of Col4a1 mRNA and protein (Fig. S1D, E). Consequently, sgRNA #4, targeting the sequences in exon 1 as shown in Fig. 1, was chosen for further use. As a control, we used sgRNA targeting the mouse *Rosa26* locus. We confirmed that lentivirus-mediated transfection of the selected sgRNA into the Cas9-expressing BECs suppressed the levels of *Col4a1* mRNA and protein in a dose-dependent manner (Fig. 1B, C).

**Figure 1.**
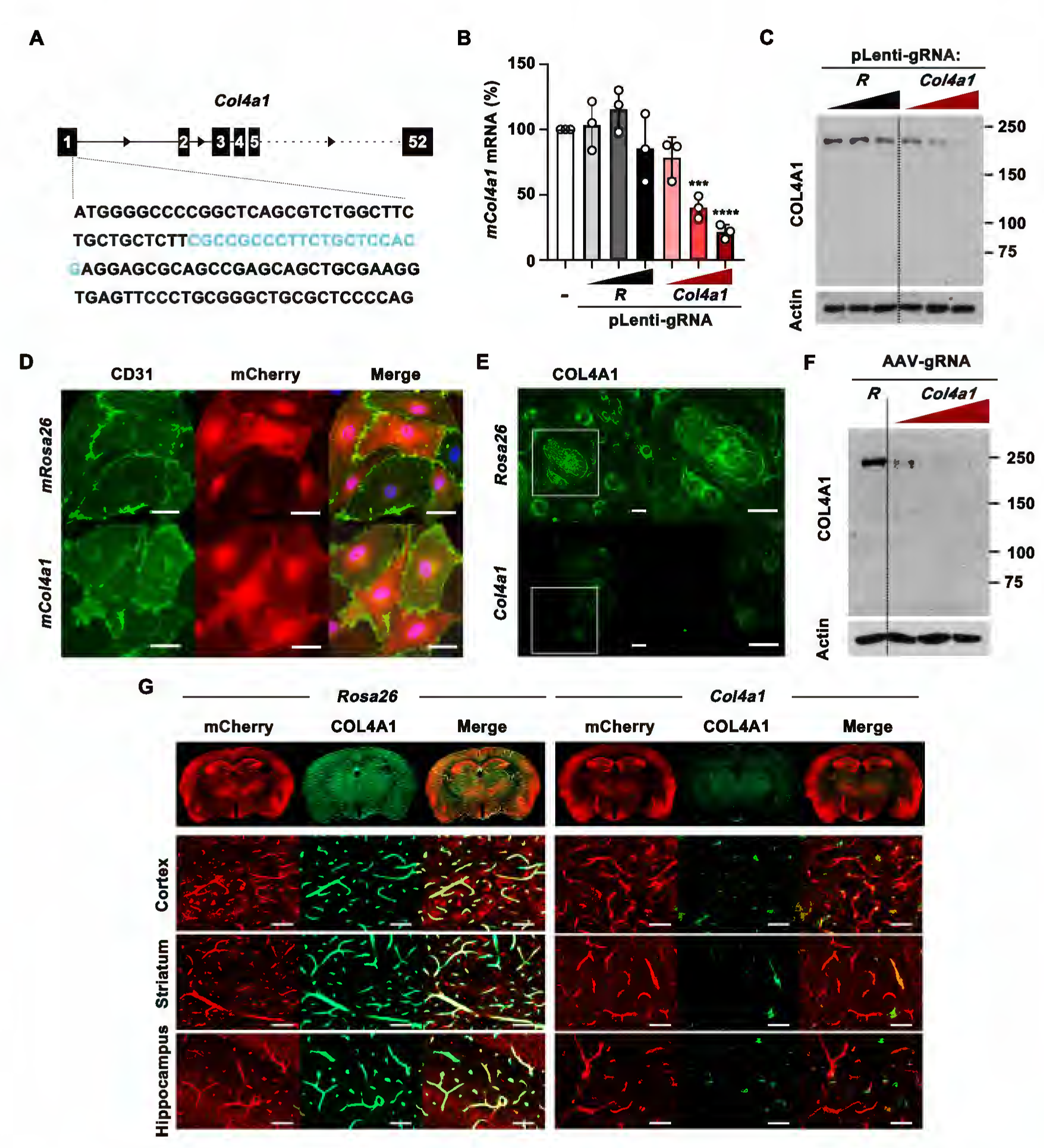
Depletion of COL4A1 in mouse brain endothelial cells by CRISPR/Cas9-mediated *Col4a1* gene editing. **(A)** A schematic diagram of the mouse *Col4a1* exon 1 targeted by the #4 sgRNA candidate. Nucleotide sequences in blue indicate the ones targeted by #4 sgRNA. **(B, C)** Quantitative RT-PCR and Western blot analysis of the primary brain endothelial cells (BECs) obtained from Cas9 transgenic mice. BECs were treated with the Lentivirus expressing sgRNAs targeting the mouse *Rosa26* locus (*R*) or *Col4a1* with increasing dosages (N = 3, \*\*\**p* < 0.001 and \*\*\*\**p* < 0.0001 compared to untreated control by one-way ANOVA followed by Tukey’s *post hoc* analysis). **(D, E)** *In vivo* editing of *Col4a1* in BECs using AAV-BR1. AAV-BR1-*Rosa26* or -*Col4a1* sgRNAs tagged with mCherry were injected into the retro-orbital sinus of 8-week-old Cas9-expressing transgenic mice. After 3 weeks, primary BECs were obtained and cultured. Cultured BECs were immunoreactive with the endothelial cell marker CD31, and the majority of CD31-positive BECs expressed mCherry. COL4A1 immunoreactivities observed in BECs from animals with AAV-BR1-*Rosa26* almost disappeared in BECs from animals with AAV-BR1-*Col4a1*. Scale bars = 20 µm. **(F)** Western blot analysis of COL4A1 protein in BECs treated with increasing dosages of AAV (1.0, 2.5, and 5.0 × 10^13^ GC/kg). Beta actin was used as an internal loading control. **(G)** Representative images of the brain tissue sections taken from animals injected with AAV-BR1-*Rosa26* or - *Col4a1*-sgRNA. Brain tissues were subjected to immunofluorescence staining using antibodies against mCherry and COL4A1. Magnified images from the cerebral cortex, striatum, and hippocampus were shown below. Scale bars = 20 µm.

For *in vivo* editing of *Col4A1* specifically in BECs, we employed AAV-BR1, a viral vector tailored for brain microvasculature endothelial cells. AAV-BR1 is modified from AAV2 and demonstrates high specificity and long-term transduction efficiency for brain ECs ^15^. To confirm its efficacy, AAV-BR1 containing GFP (AAV-BR1-GFP) was administered to 8-week-old mice via a retro-orbital sinus route. The GFP signals were colocalized with CD31+ brain ECs (Fig. S2A). More than 90% of CD31 positive vessels were colocalized with GFP signals, indicating that most of the brain microvessels were transduced by the AAV-BR1 viral vector. In contrast, GFP signals were not localized with astrocytes or microglial cells. Consistent with the previous report ^15^, scattered neuron transduction was observed within the cerebral cortex (Fig. S2A). However, the frequency of neuronal transduction was extremely low compared to that of endothelial transduction. While the GFP signals were robustly observed in the brain, they were rarely detected in the peripheral organs, such as the liver, lung, and heart (Fig. S2B).

The sgRNA #4 sequence was packaged into AAV-BR1 vector with CAG promoter driven mCherry to facilitate tracing (hereafter referred to as “AAV-BR1-*Col4a1*-sgRNA”), and the AAV-BR1-*Col4a1* sgRNA was injected via a retro-orbital route into adult Cas9-expressing transgenic mice. Three weeks after the injection, primary BECs were harvested from the animals and cultured for 10 days. The *ex vivo* cultured BECs were positive for the endothelial cell marker CD31, and most CD31-positive endothelial cells expressed mCherry tagged at the 3’ end of sgRNA sequence, indicating high BEC transduction efficiency of AAV-BR1-*Col4a1*-sgRNA (Fig. 1D). Immunofluorescence staining revealed that the *ex vivo* cultured BECs from animals with AAV-BR1-*Col4a1*-sgRNA expressed a very low level of COL4A1 compared to those from animals with control AAV-BR1-*Rosa*-sgRNA (Fig. 1E). Western blot analysis confirmed the almost complete disappearance of COL4A1 protein in cultured BECs from animals treated with AAV-BR1-*Col4a1*-sgRNA at a dosage of 0.5 × 10^13^ GC/ml or higher (Fig. 1F). In the brain, collagen IV (COL4A1) was found to localize specifically around blood vessels, similar to the CD31 immunoreactivities of (Fig. 1G). In animals injected with AAV-BR1-*Rosa*-sgRNA, most collagen IV-containing blood vessels were colocalized with mCherry signals. Injection of AAV-BR1-*Col4a1*-sgRNA resulted in a substantial reduction of collagen IV immunoreactivity (Fig. 1G), indicating the effectiveness of our *Col4A1* gene editing approach.

### Col4a1 depletion in BECs induces CMBs on MRI

We used 9.4T small animal MRI system to monitor potential pathological changes in the brain. Adult animals (8 weeks old) were injected with AAVs containing sgRNA targeting either *Rosa26* or *Col4a1*. Three months after the AAV-BR1-*Col4a1*-sgRNA injection, we observed small and rounded hypointense spot-like structures scattered in the cortex and hippocampus in T2-weighted MRI (Fig. 2A). At this time point, CMBs were rarely observed in subcortical structures such as the striatum, thalamus, and brainstem (Fig. 2A, B). However, the number of CMBs on MRI sharply increased at 6 months, especially in the cortex and striatum. Additionally, CMBs were more frequently observed in the thalamus, hypothalamus, and brainstem. The preferential distribution of CMBs in cortical regions and deep gray matter structures appeared to be consistent with common sites of microbleed occurrence in humans ^16–18^. No CMB-like signals were found in animals injected with AAV-BR1-*Rosa26*-sgRNA, indicating that the CMB-like lesions were not caused by the AAV-BR1 virus injection. Importantly, no animals exhibited macro-hypointensity lesions with obvious tissue damage in nearby areas, suggesting that no ICHs occurred.

**Figure 2.**
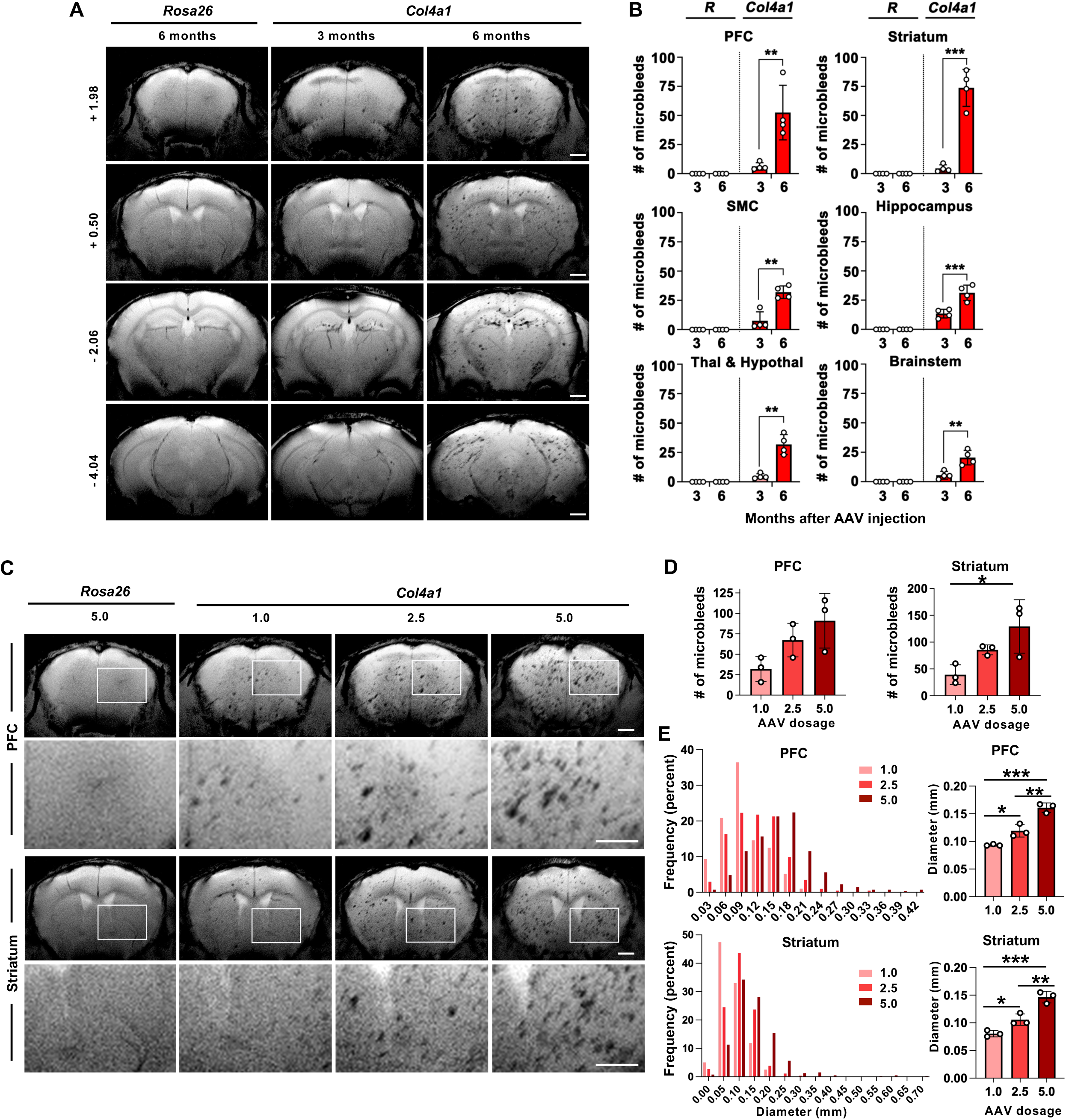
Col4A1 depletion in BECs induces cerebral microbleeds on magnetic resonance imaging. **(A)** Representative T2-weighted magnetic resonance images (MRIs) displaying cerebral microbleeds (CMBs) at 3 or 6 months after injection with AAV-BR1-*Rosa26* (*R*) or -*Col4a1*-sgRNA. The selected coronal images were obtained at the specified distances from the bregma, as indicated on the left. Scale bars = 0.5 mm. **(B)** Quantitative analysis showing the number of CMBs counted on T2-weighted images at 3 or 6 months after the AAV-BR1-*Rosa26* or -*Col4a1*-sgRNA injection at a dosage of 2.5 × 10^13^ GC/kg. PFC: Prefrontal cortex, SMC: Sensory Motor Cortex, Thal: Thalamus, and Hypothal: Hypothalamus. N = 4 animals per group. (** *p*< 0.01 and *** *p* < 0.001 by Student’s t-test). Error bars represent the SEM. **(C)** Representative T2-weighted MRI obtained 6 months after the AAV-BR1-*mRosa26* (*R*) or -*mCol4a1*-sgRNA (*Col4a1*) injection at three different doses (1.0, 2.5, and 5.0 × 10^13^ GC/kg). Scale bars = 0.5 mm. **(D)** Quantitative analysis showing the number of CMBs in the PFC and striatum 6 months after AAV injection at three different doses (1.0, 2.5, and 5.0 10^13^ × GC/gm). Each circle represents an individual animal subject. * *p* < 0.05 by one-way ANOVA followed by Tukey’s *post hoc* analysis. N = 3 animals per condition. Error bars represent the standard deviation. (E) Frequency histograms of CMB diameters in the PFC and striatum (left) and quantitative graphs of comparing the mean diameter of CMBs in the PFC and striatum (right). Each circle represents the mean CMB diameter of an individual animal subject. * *p* < 0.05, ** *p* < 0.01, and ** *p* < 0.001 by one-way ANOVA followed by Tukey’s *post hoc* analysis. For the frequency histograms, N = 96, 202, and 268 CMBs for 1.0, 2.5, and 5.0 × 10^13^ dosage groups, respectively. For the bar graphs, N = 3 animals per condition. Error bars represent the standard deviation.

We also investigated whether the dosage of AAV-delivered sgRNA would influence the severity of CMBs. We injected animals with three different doses of AAV-BR1-*mCol4a1*-sgRNA: 1.0, 2.5, and 5.0 × 10^13^ GC/kg. We then counted the number of CMBs 6 months after in the prefrontal cortex and striatum, where CMBs were most frequently found. There was a clear tendency for a dose-dependent increase in the number of CMBs on MRI in both regions, although the increases were statistically significant only in the striatum. (Fig. 2C, D). In animals receiving the highest dose, more than 100 CMBs were observed on a single MRI plane, with almost the entire prefrontal cortex and the striatum studded with CMBs of varying sizes. Most CMBs were less than 300 μm in diameter, with an average diameter of 150 μm or less in all three dosage groups (Fig. 2E). CMBs larger than 500 μm were almost exclusively found in animals receiving the highest dose, with the largest CMB measuring approximately 700 μm in diameter in this group. Despite the large paramagnetic signals in this group, there was no evidence of macroscopic hemorrhage accompanied by changes in signal intensity in the parenchymal regions adjacent to the CMBs. In contrast, the majority of CMBs in animals receiving the lowest dosage were less than 100 μm. When the frequency distribution of CMB size was compared, the differences in size distribution were statistically significant among all dosage pairs (*p* < 0.001 by Kolmogorov-Smirnov test) (Fig. 2E). The mean diameter of CMBs was also significantly different among all the different dosage groups (Fig. 2E), demonstrating a dose-dependent influence of AAV dosages on the CMB size.

### Intravital imaging and histological evaluation

To visualize the CMBs in the living brain, we used intravital imaging with a two-photon microscope in Cas9 transgenic animals 6 months after AAV injection. A cranial window was created over the sensorimotor cortex, and FITC-conjugated dextran (150 kDa) was intravenously injected to visualize the cerebral blood vessels. Immediately after the FITC-dextran injection, we were able to visualize the fluorescent dye leaking from small cerebral blood vessels in an animal with AAV-BR1-*Col4a1*-sgRNA (Fig. 3A, Movie S1), indicating the presence of structural disintegration in the blood vessels. Three-dimensional projection images clearly showed varying levels of extravasation of fluorescently labeled dextran (Fig. 3B, Movie S2). In contrast, there was no evidence of extravasation in animals that received the control AAV-BR1-*Rosa*-sgRNA injection.

**Figure 3.**
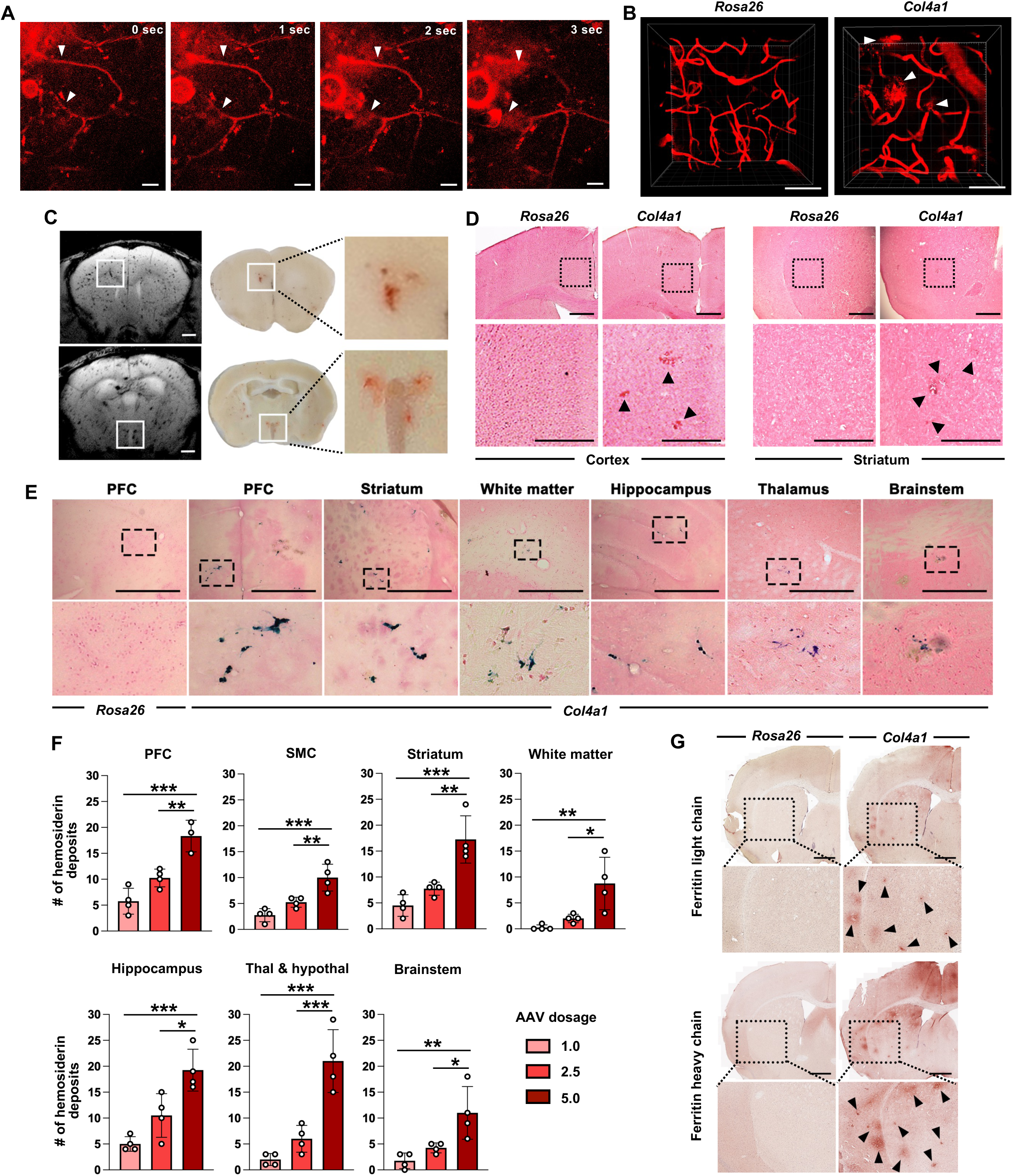
Intravital imaging and histological evaluation of CMBs. (A) Time-lapse images of cerebral blood vessels obtained with one second interval by a two-photon microscope. The intravital imaging was performed 6 months after AAV injection through a cranial window installed over the sensorimotor cortex. The brain vessels were labeled by injecting 100 µL of 150 kDa FITC-dextran via the retro-orbital route. White arrowheads indicate the areas of the small vessels where the FITC-dextran is leaking out. Scale bars = 20 μm. (B) Representative two-photon 3D projection images visualizing cerebral microvessels. White arrows indicate extravasation of fluorescently labeled dextran in an animal injected with AAV-BR1-*Col4a1*-sgRNA. There was no evidence of extravasation in an animal with AAV-BR1-*Rosa26*-sgRNA. Scale bars: 50 μm (C) Correlation of paramagnetic signals indicating CMBs using T2* MRI with corresponding brain slices showing hemorrhagic signs in mice with AAV-BR1-*Col4a1*-sgRNA injection. Scale bars = 1 mm. (D) Representative images of the cerebral cortex and striatum stained with hematoxylin and eosin. The brain tissues were obtained from animals 6 months after AAV injection. The arrowheads indicate presumed microbleeding. Scale bars = 0.5 mm. (E) Representative images of hemosiderin deposits visualized by Prussian blue staining. The boxed regions in the upper panels are magnified in the lower panels. Scale bars = 0.5 mm. (F) Quantitative analysis of hemosiderin deposits 6 months after the AAV-BR1-*Col4a1*-sgRNA injection at varying dosages (1.0, 2.5, and 5.0 × 10^13^ GC/kg). N = 4 animals for each dose of AAV injection. *, **, and *** indicate *p* < 0.05, *p* < 0.01, and *p* < 0.001 by one-way ANOVA followed by Tukey’s *post hoc* analysis. Error bars represent the standard deviation. (G) Representative images of brain slices obtained from AAV-*Rosa26*-sgRNA or AAV-*Col4a1*-sgRNA-injected adult mice. Immunohistochemical staining was performed using antibodies against ferritin light (top panel) and heavy chains (bottom panel). The boxed regions in the upper panels are magnified in the lower panels. Black arrowheads indicate the immunoreactivities of ferritin light and heavy chains. Scale bars = 1 mm.

Histological examination was conducted 6 months after AAV injection. CMBs were clearly visible on the coronal sections of the brain tissue that had been fixed with paraformaldehyde (Fig. 3C). However, the density of CMBs visually identified on the tissue was lower than that detected on the MRI. Several CMBs appeared pinkish or dark reddish in color, suggesting that the visually identified CMBs may be more recent ones. Certain CMBs were traced on corresponding MR images, enabling anatomical and radiological correlations (Fig. 3C, boxed regions). In coronal brain sections stained with hematoxylin and eosin, scattered dark brown or golden-brown spots were observed throughout the brains of animals injected with AAV-BR1-*Col4a1*-sgRNA (Fig. 3D). These spots indicate microscopic evidence of hemosiderin deposition ^19^. To visualize iron deposition in the brain tissue more directly, Prussian blue staining was performed. This staining revealed dark blue deposits of various sizes in regions where CMBs were frequently identified on MRI scans (Fig. 3E). The number of the Prussian blue-positive spots seemed to be higher than those found with hematoxylin and eosin staining. We quantified the number of Prussian blue-stained deposits in various regions of the brain from animals with varying doses of AAV-BR1-*Col4a1*-sgRNA (Fig. 3F). Overall, there was a correlation between the regions where CMBs were frequently found on MRI and histological sections. The number of CMBs on histological sections was also dependent on the viral dosages in both evaluations. We also performed immunohistochemical staining with antibodies against ferritin light and heavy chains (Fig. 3G). The cortex and striatum showed both light and heavy chain ferritin immunoreactivities, and these immunoreactive areas were significantly larger than the hemosiderin spots observed with H & E or Prussian blue-stained iron deposits. This indicates that ferritin levels may increase in areas near substantial hemosiderin or iron deposits.

We examined if targeted *Col4a1* deletion induced CVSD pathologies other than CMBs, particularly ischemic changes. There was no single incidence of lacunar or microinfarctions in animals injected with either control AAV or AAV-BR1-*Col4a1*-sgRNA (Fig. S3A). Importantly, we did not observe any changes in the integrity of white matter in the corpus callosum as shown by myelin-specific Eriochrome cyanine RC staining (Fig. S3A). Furthermore, immunoreactivity against MBP in the corpus callosum did not differ between the two groups (Fig. S3B), indicating that the targeted deletion of *Col4a1* in the brain microvessels does not result in ischemic white matter injury.

### Activation of glial cells induced by CMBs

We investigated whether hemosiderin deposits can lead to degeneration of neural cells adjacent to microbleeding sites. There was no change in the density of NeuN-positive neurons in the cortical regions close to Prussian blue-stained hemosiderin deposits (Fig. S4A). Immunohistochemical staining of MBP also did not reveal any signs of degeneration of myelinated fibers passing through or adjacent to the hemosiderin deposits (Fig. S4B). Next, we examined alterations in glial activation resulting from iron deposition. GFAP staining revealed reactive astrocytes located near the Prussian blue-positive iron deposits (Fig. 4A). Additionally, we noted an increased intensity of IBA-1 immunoreactivity, indicating activation of microglial cells. Notably, the IBA-1 immunoreactivity seemed to encircle the iron deposits (Fig. 4B), suggesting that microglial cells may be phagocytosing the iron spots. We also found that the iron deposits were sometimes incorporated into CD45-positive cells (Fig. 4C), indicating that blood-derived macrophages were also recruited to the CMBs.

**Figure 4.**
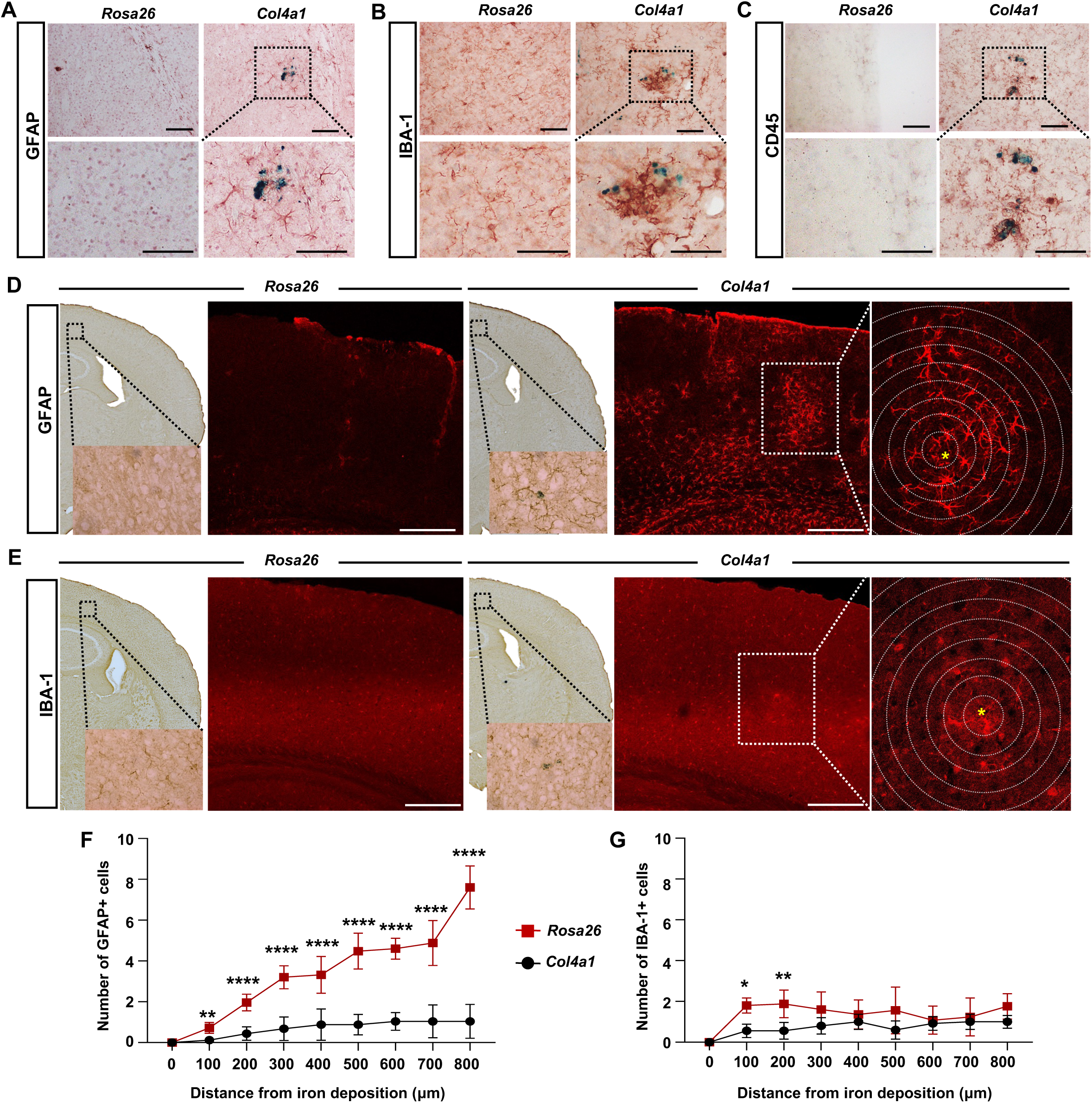
Activation of glial cells induced by CMBs (A–C) Representative immunohistochemical images of the brain slices obtained from AAV-*Rosa26* or AAV-*Col4a1*-injected mice. The hemosiderin-laden cells in the brain slices were stained with antibodies against GFAP, IBA1, and CD45, followed by staining using the Prussian blue reaction method. Scale bars = 20 μm. **(D, E)** Representative low-magnification images of the brain slices immunostained with GFAP (D) and IBA-1 (E) followed by the Prussian blue staining. The spatial distribution of iron deposits was visualized in brain sections stained with the Alexa Fluor 594, as the dark field image provided higher contrast. Scale bars = 500 μm. **(F, G)** Quantification graphs comparing the number of astrocytes (F) and microglial cells (G) contacted by concentric circles evenly spaced from the iron deposit. *, **, and **** indicate *p* < 0.05, *p* < 0.01, and *p* < 0.0001 by repeated measures two-way ANOVA followed by Bonferroni *post hoc* analysis. Error bars represent the SEM.

It is noteworthy that the spatial pattern of astrocytic activation differed significantly from that of IBA-1 positive microglial cells. GFAP immunoreactivity was observed not only in the vicinity of an iron deposit but also at considerable distances from it. Occasionally, the far-reaching GFAP immunoreactivities overlapped with areas surrounding different iron spots, leading to extensive activation of astrocytes (Fig. 4D). In contrast, microglial activation appeared to be limited to cells that were either in close proximity to or actively phagocytosing the iron deposits (Fig. 4E). We measured the number of astrocytes and microglial cells contacted by concentric circles evenly spaced from the iron spot. We found that the number of astrocytes increased with greater distance (Fig. 4F), while the number of microglial cells remained unchanged beyond 100 to 200 μm (Fig. 4G).

### Ultrastructural analysis of brain microvessels

To further examine structural changes in the brain microvessels, we used correlative light and electron microscopy (CLEM) which combines fluorescence images with the electron microscopic (EM) features of cerebral microvessels. In control mice that were injected with AAV-BR1-*Rosa26*-sgRNA, CD31-positive endothelial cells rarely came into contact with IBA1-positive microglial cells (Fig. 5A). However, in animals with AAV-BR1-*Col4a1*-sgRNA, microglial cells surrounded the BECs and were associated with hemosiderin deposits near the blood vessels (Fig. 5B). It appeared that several microglial cells were recruited to areas with electron-dense hemosiderin deposits and contained these deposits within distinct microglial plasma membranes (Fig. 5B).

**Figure 5.**
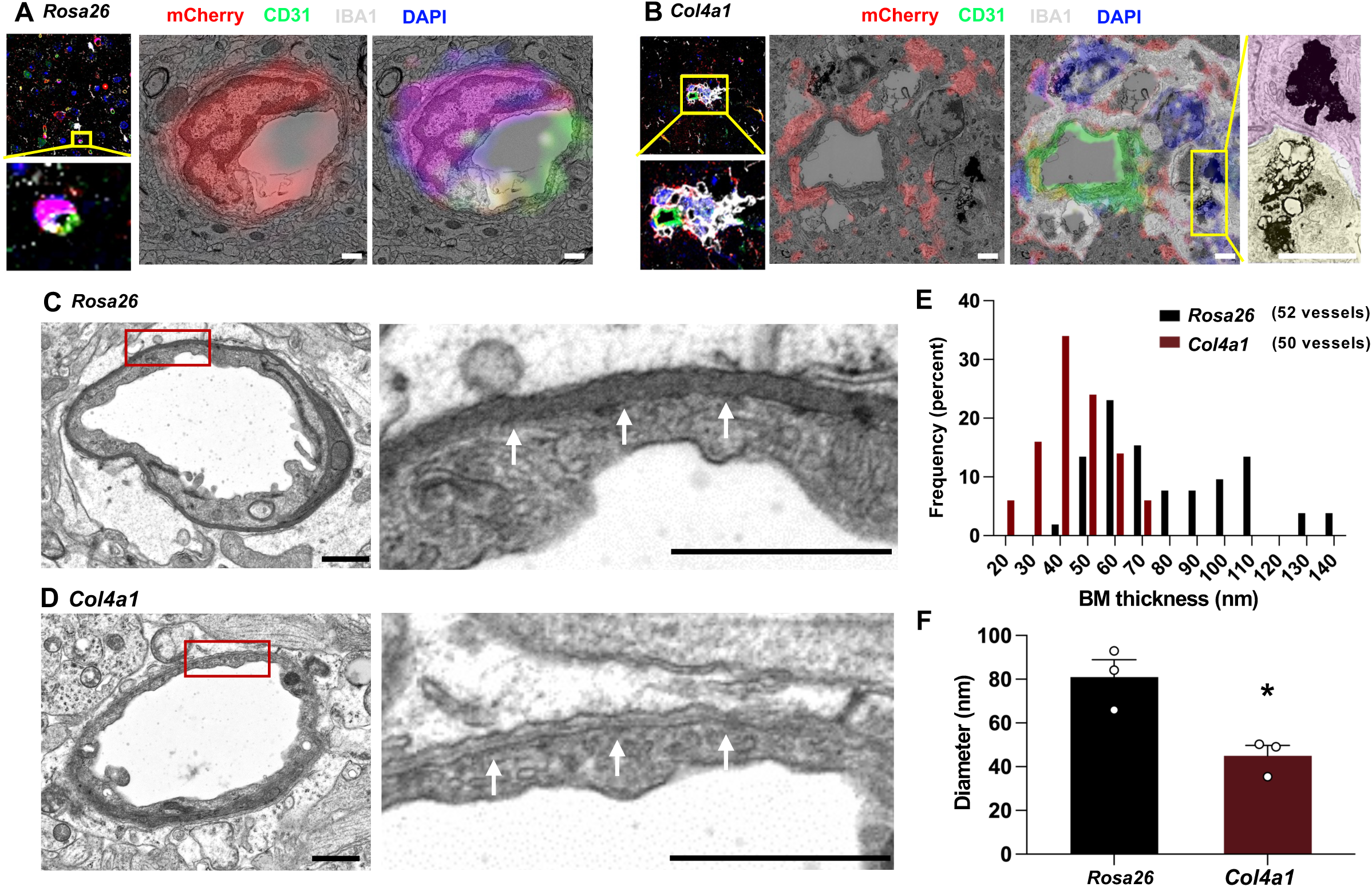
Ultrastructural analysis of brain microvessels (A, **B)** Representative correlative light and electron microscopy (CLEM) images of brain microvessels obtained from AAV-BR1*-Rosa26*-sgRNA **(A)** or AAV-BR1-*Col4a1*-sgRNA **(B)** injected mice. Brain endothelial cells were visualized by CD31 immunoreactivity (green) combined with mCherry signals, while microglial cells were stained using an antibody against IBA1 (white). The boxed regions (yellow square boxes) on the left are magnified below, and electron-dense deposits in the yellow rectangular box (B) are shown in greater detail on the right. Different microglial cells have been colored differently in the magnified image. Scale bars = 1 μm. **(C, D)** Representative electron microscopy (EM) images of microvessels in coronal brain sections from mice injected with AAV-BR1-*Rosa26*-sgRNA **(C)** and AAV-BR1-*Col4a1*-sgRNA **(D)**, respectively. The rectangular boxed areas on the left are magnified on the right. Scale bars = 1 μm. **(E)** A frequency histogram of BM thickness. To quantify the thickness of the BM, three lines were drawn perpendicular to the outer boundaries of the BM in each microvessel, and their lengths were measured using image analysis software. A total of 52 and 50 vessels were analyzed for the control and Col4a1 groups, respectively. **(F)** A quantification graph of the average BM thickness. * indicates *p* < 0.05 as determined by an unpaired Student’s t-test. N = 3 animals per group.

To evaluate the consequence of *Col4a1* disruption on the BM structures, we conducted conventional EM analysis of cerebral microvessels in brain slices that were injected with AAV. Our observations revealed that the thickness of the BM in animals treated with AAV-BR1-*Col4a1*-sgRNA was significantly reduced compared to that in control animals (Fig. 5C, D). Additionally, we noted a decrease in the electron density within the thinned BM in the Col4a1 gene-edited animals. We measured the BM thickness of 52 vessels from three control animals and 50 vessels from three animals with Col4a1 editing that were not accompanied by hemosiderin depositions. We then compared the frequency distribution of BM thickness between the two groups (Fig. 5E). Our analysis revealed that a higher percentage of vessels in the control group had BM structures thicker than 60 nm. In contrast, over 50% of the vessels in the AAV-BR1-Col4a1-sgRNA group exhibited BM thicknesses of 40 nm or less. The Kolmogorov-Smirnov test indicated a significant difference in the frequency distribution between the two groups (*p* < 0.001). Furthermore, the mean BM thickness was significantly smaller in the AAV-BR1-Col4a1-sgRNA group, as demonstrated by an unpaired Student’s t-test (Fig. 5F). Thus, gene editing of Col4a1 in adult brain microvessels led to a structural disintegration of the BM structures in unruptured brain microvessels.

### Cognitive decline and motor incoordination by *Col4a1* deletion in brain ECs

To assess whether the presence of multiple CMBs due to targeted *Col4a1* deletion was linked to neurological impairments, the animals underwent a series of behavior tests, including the Novel object recognition (NOR), Y-maze, and Rotarod test. (Fig. 6A, E, I). In the NOR test, animals with *Col4a1*-targeting sgRNA began to show a decrease in discrimination index as early as 2 months after AAV injection (Fig. 6B). Their inability to discriminate a novel object declined even further through 6 months. Repeated measures two-way ANOVA revealed significant inter-group differences between the two groups (*F_(1, 30)_* = 75.95, *p* < 0.0001). When the data were analyzed separately based on the gender of animals, both sexes showed a similar decrease in the discrimination index (male: *F_(1, 14)_* = 33.79, *p* < 0.0001, female: *F_(1, 14)_* = 41.86, *p* < 0.0001). Female mice appeared to exhibit the deficit earlier than males, but the extent of the deficits at the last time point was comparable between the sexes (Fig. 6C, D). Y-maze testing for short-term executive memory showed comparable performance until 4 weeks after AAV injection (Fig. 6F). At 5 months post AAV administration, mice treated with AAV-BR1-*Col4a1*-sgRNA mice demonstrated decreased alternation triplet percentage, becoming more pronounced at 6 months (repeated measures two-way ANOVA: *F_(1, 30)_* = 5.481, *p* < 0.05). Both sexes showed a similar trend, but the decline at 5 and 6 months was more noticeable in males, resulting in statistical significance only in male mice (male: *F_(1, 14)_* = 6.09, *p* < 0.05, female: *F_(1, 14)_* = 1.47, *p* = 0.228) (Fig. 6G, H). The rotarod test is commonly used to evaluate motor coordination in rodents ^20^. Mice with control sgRNA showed a slight decrease in their ability to remain on the rotarod as they aged (Fig. 6J). However, mice with *Col4a1*-targeting sgRNA exhibited a rapid decline from 3 months after injection, revealing a significant difference (*F_(1, 30)_* = 17.87, *p* < 0.001). The difference was statistically significant in both male and female mice (male: *F_(1, 14)_* = 16.65, *p* < 0.01, female: *F_(1, 14)_* = 4.83, *p* < 0.05), with the motor coordination deficit being more noticeable in males (Fig. 6K, L). Post-hoc Bonferroni tests revealed significant differences from 3 to 6 months in males but only at 6 months in females. General locomotor activity showed no significant differences between groups (Fig. S5A-C), nor did body weight changes (Fig. S5D-F), excluding potential influences of differences in motivation or systemic factors on motor activities during all three tests.

**Figure 6.**
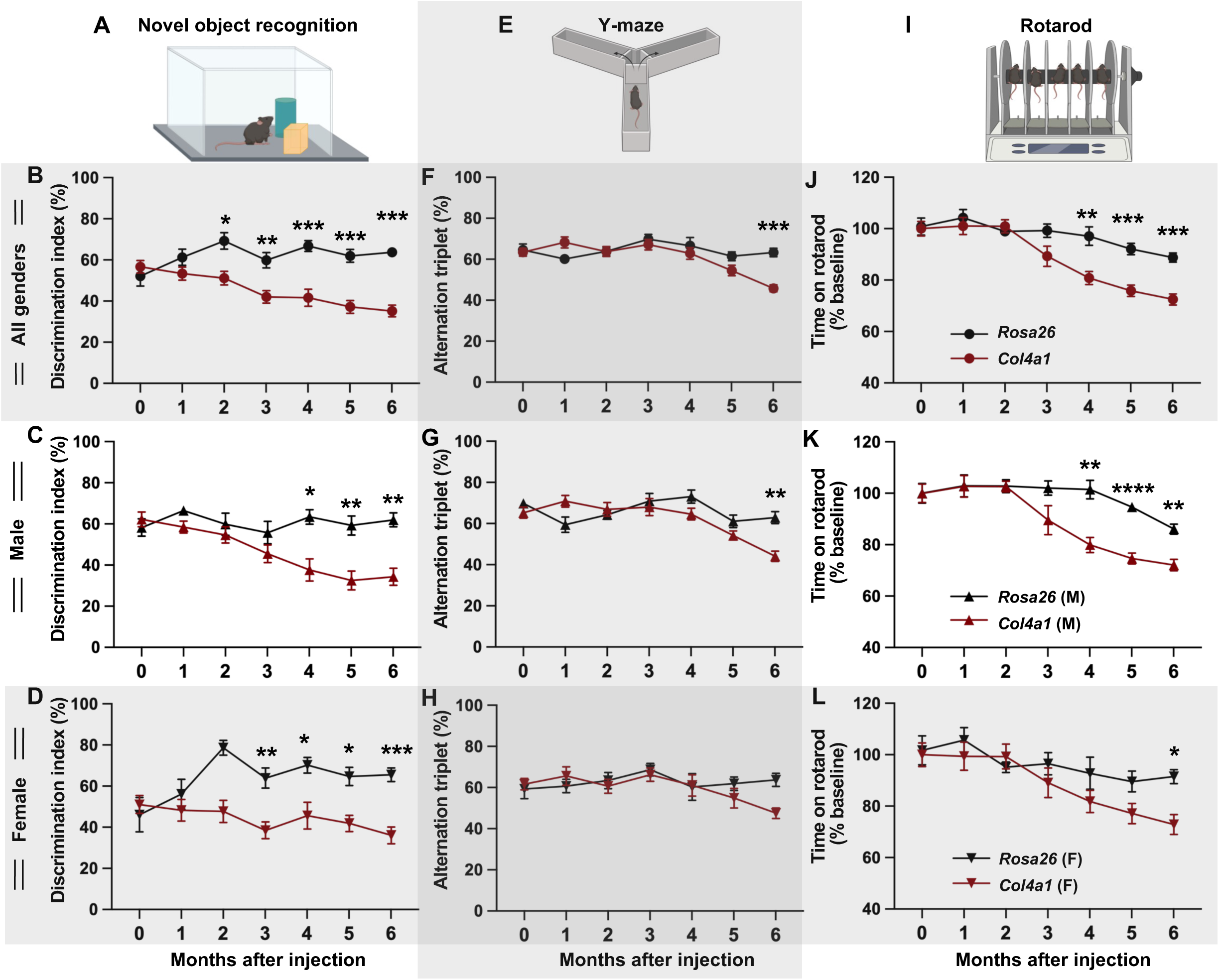
Cognitive decline and motor incoordination by Col4a1 depletion in brain microvessels (A-D) Novel object recognition test. **(A)** A schematic diagram of the novel object recognition test. Animals were injected at the age of 8 weeks. The test was conducted before the injection and then once a month for six months following the injection. The discrimination index was calculated from all animals, regardless of gender **(B)**, as well as separately for male **(C)** and female **(D)** animals. The discrimination index was defined as the time spent exploring a new subject divided by the time spent exploring both objects. **(E-H)** Y-maze test. **(E)** A schematic diagram of the Y-maze test. The percent alternation triplet was obtained from all animals, regardless of gender **(F)**, as well as separately for male **(G)** and female **(H)** animals. An alternation triplet was considered if there was a consecutive sequence of entries into all three arms. The formula to calculate the percent alternation triplet is described in the Methods section. **(I-L)** (I) A schematic diagram of the rotarod test. The amount of time taken until animals fell from the apparatus was recorded from all animals, regardless of gender **(J)**, as well as separately for male **(K)** and female **(L)** animals. * *p* < 0.05, ** *p* < 0.01, and *** *p* < 0.001 by repeated measure two-way ANOVA followed by post-hoc Bonferroni test. N = 14 (7 per gender) and 18 (9 per gender) for control sgRNA against Rosa26 and targeting Col4a1, respectively.

### Association of single nucleotide polymorphisms (SNPs) in genes related to collagen IV with CMBs

The results of the above animal study suggest that alterations in collagen IV of the BM in the brain microvessels may play a significant role in the development of CMBs in human patients. We hypothesized that human *COL4A1* gene variants might influence CMB susceptibility in aging populations. We analyzed data from 836 participants with available T2*-weighted gradient echo (GRE) or susceptibility-weighted imaging (SWI) MRI sequences from the BICWALZS biobank initiative, a Korean government-supported multi-center project designed to facilitate research using biospecimens and clinical data from elderly subjects with subjective memory impairment ^21^. Following previously reported criteria ^22,23^, study subjects were stratified into three groups based on CMB burden: no CMBs (Group I, N = 607, 72.6%), 1-4 CMBs (Group II, N = 173, 20.7%), and ≥ 5 CMBs (Group III, N = 56, 6.7%) (Table 1). Representative MR images demonstrating the spectrum of CMB pathology across these groups are presented in Fig. S6. Statistical analysis revealed significant differences in the mean age and K-MMSE score across the groups (*p* < 0.05), with Group II showing significantly advanced age and lower cognitive performance compared to Group I (Table 1). Chi-square analysis of vascular risk factors showed significantly higher prevalence of hypertension and prior stroke history in both Groups II and III relative to Group I (*p* < 0.05 for both comparisons), while diabetes and dyslipidemia rates were comparable across the three groups.

**Table 1.**
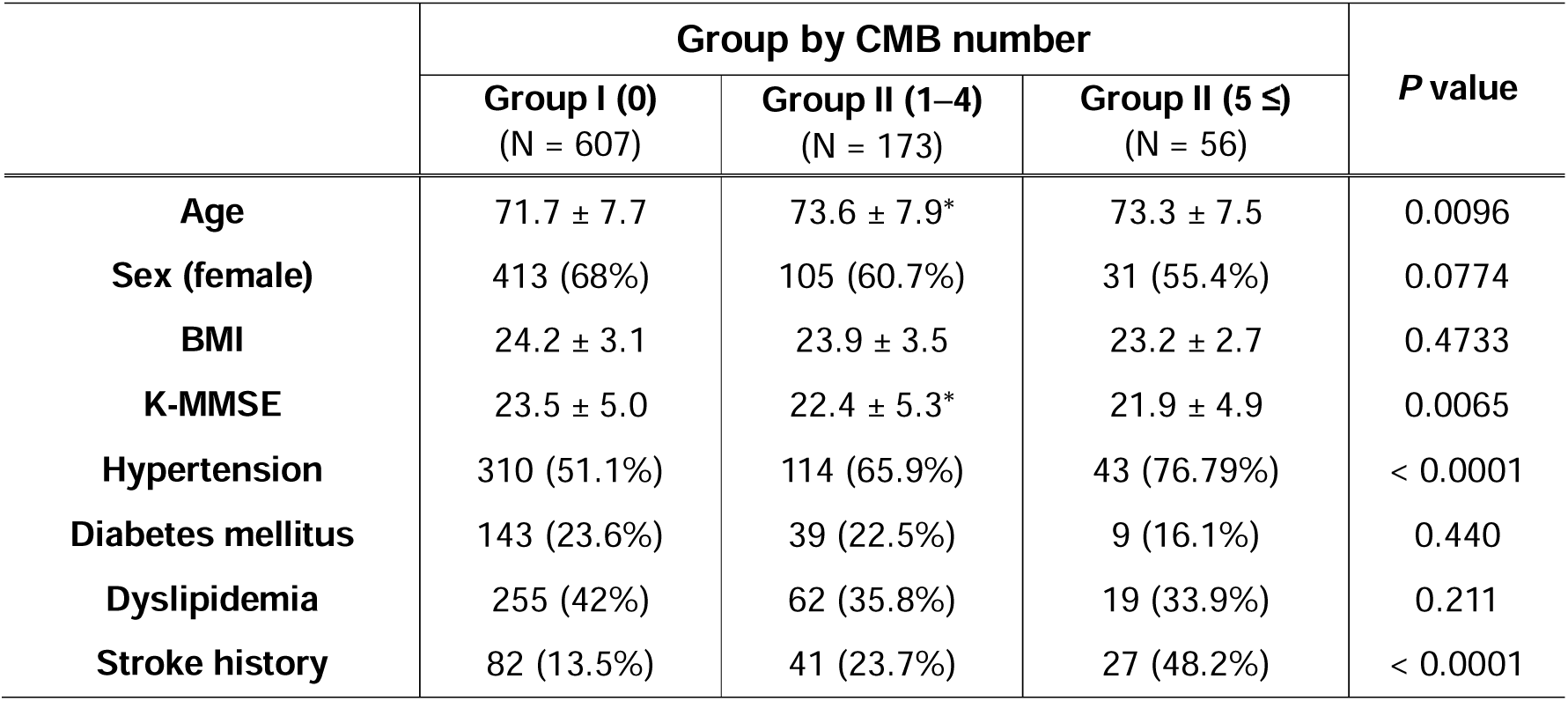
Demographic information and clinical characteristics of the subjects included. Continuous variables such as age, BMI, and K-MMSE (Korean mini-mental status examination) were expressed as average values with standard deviation. Categorical variables such as sex, vascular risk factors, and stroke history were shown as the number (%) of subjects with the corresponding condition. P values were calculated by one-way ANOVA for continuous variables and chi-square contingency test for categorical variables. * *p* < 0.05 compared to Group 1 by one-way ANOVA followed by Tukey’s *post hoc* analysis.

We investigated the relationship between genetic variants in the *COL4A1* gene and the CMB burden observed in MRI scans using linear regression analysis. Among the 84 SNPs identified in the study cohort, only rs3825469 demonstrated a statistically significant association with the number of CMBs, achieving a *p*-value below 0.0006, the threshold adjusted for multiple comparisons using Bonferroni correction (Table S1). To identify SNPs associated with the CMB burden in genes other than *COL4A1*, we conducted a genome-wide association study using linear regression modeling. No polymorphisms reached genome-wide significance at the conventional suggestive threshold (*p* < 5 × 10[^5^) (Fig. S7). We subsequently conducted pathway analysis on the top 100 genes, which included 107 SNPs, ranked by ascending *p*-value (see Table S2), and constructed a protein-protein interaction network that incorporated COL4A1. This network analysis revealed several hub proteins with multiple interactions, among which COL4A1 exhibited the most extensive connectivity (Fig. 7A). We quantified the strength of the interactions between COL4A1 and its interacting partner proteins and found that matrix metalloproteinase 2 (MMP2) has the strongest interaction with COL4A1 (Fig. 7B). Importantly, tissue inhibitor of matrix metalloproteinase 2 (TIMP2), a specific endogenous inhibitor of MMP2 activity ^24^, demonstrated robust connectivity with both MMP2 and COL4A1 (Fig. 7A, B). Targeted regression analysis found no significant *MMP2* SNPs after multiple testing correction (Table S3). However, we identified four *TIMP2* SNPs significantly associated with CMB number, all achieving *p*-values below 0.0022, the Bonferroni-corrected significance threshold (Table S4).

**Figure 7.**
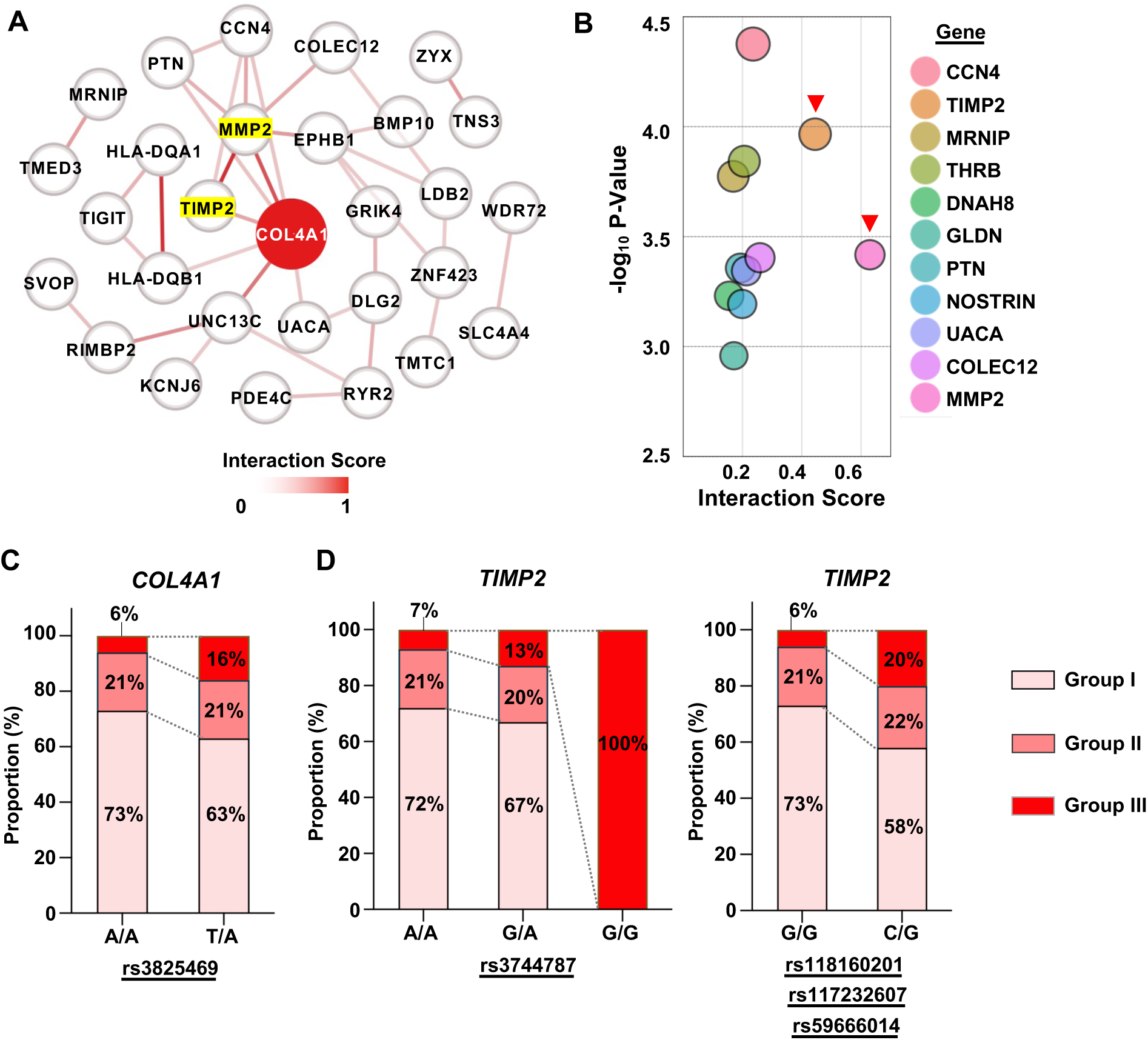
Protein-protein interaction analysis and the graphical representation of the contingency table analysis. **(A)** Generation of a protein-protein interaction map using STRING analysis. The top 100 genes ranked by ascending *p*-values obtained from a genome-wide association study (GWAS) were used for the analysis. The strength of nodal interactions is color-coded. **(B)** A bubble plot presentation of the interaction scores of the genes that interact with *Col4a1*. The Y-axis represents *p*-values of the genes obtained from the GWAS linear regression. MMP2 and TIMP2 are highlighted by arrowheads. **(C, D)** Graphical representations of the contingency table analysis of genotype distribution of **(C)** *Col4a1* and **(D)** *TIMP2* variants across the trichotomized CMB severity groups (Group I, II, and III).

To characterize how these five significantly associated SNPs (one in *COL4A1* and four in *TIMP2*) contribute to CMB formation susceptibility, we analyzed genotype distributions across our trichotomized CMB severity groups using contingency table analysis (Table 2). For the *COL4A1* rs3825469 variant, we found that comparable frequency of heterozygous (T/A) and wild-type (A/A) genotypes in Group II, but Group I had fewer heterozygotes relative to major allele homozygotes (63% vs. 73%), while Group III showed an increase in heterozygote representation (16% vs. 6%) (Table 2, Fig. 7C). However, chi-square analysis did not reach statistical significance for genotypic distribution differences across CMB severity groups (χ² = 3.660) (Table 2). In contrast, the *TIMP2* variants demonstrated significant associations with CMB burden in the contingency analysis. For rs3744787 SNP, the proportion of subjects with the heterozygous genotype (G/A) was lower than that of the wild-type homozygotes (A/A) in Group 1 and higher in Group III, revealing an increased odds ratio for having a higher CMB burden (Table 2, Fig. 7D). Notably, two subjects with the homozygous variant minor allele genotype (G/G) exhibited severe microbleed pathology (≥ 5 CMBs), indicating a substantial gene dose-dependent effect on CMB susceptibility. The remaining three *TIMP2* SNPs (rs118160201, rs117232607, and rs59666014) showed identical distribution patterns, with heterozygous carriers demonstrating a nearly two-fold increased risk for severe CMB burden (OR = 1.96) compared to major allele homozygotes (Table 2, Fig. 7D). The identical contingency tables for these three variants indicated transmission as a linkage disequilibrium block, confirmed by regional plots showing their close genomic proximity and highly significant associations with each other within the *TIMP2* gene (Fig. S8). Chi-square analysis revealed significant differences in genotype distribution across CMB groups for all TIMP2 variants (χ² = 3.966 for rs3744787; χ² = 7.972 for the other linked SNPs), collectively supporting the potential mechanistic importance of *TIMP2* in CMB pathophysiology.

**Table 2.** Distribution of Snotypes across the three groups categorized by CMB number.

| Gene | SNP | Genotype | Subject number (proportion) |  |  | Subject number (Total) | OR (95% CI) | P value |
| --- | --- | --- | --- | --- | --- | --- | --- | --- |
|  |  |  | Group I | Group II | Group III |  |  |  |
| COL4A1 | rs3825469 | A/A | 525 (73%) | 151 (21%) | 40 (6%) | 716 | 1.59<br>(0.98-2.58) | 0.05575 |
|  |  | T/A | 47 (63%) | 16 (21%) | 12 (16%) | 75 |  |  |
|  |  | T/T | 0 | 0 | 0 | 0 |  |  |
| TIMP2 | rs3744787 | A/A | 505 (72%) | 146 (21%) | 36 (7%) | 687 | 1.50<br>(1.00-2.25) | 0.04644 |
|  |  | G/A | 72 (67%) | 22 (20%) | 14 (13%) | 108 |  |  |
|  |  | G/G | 0 | 0 | 2 (100%) | 2 |  |  |
|  | rs118160201 | G/G | 535 (74%) | 152 (21%) | 37 (5%) | 724 | 1.96<br>(1.22-3.16) | 0.00475 |
|  |  | C/G | 43 (58%) | 16 (22%) | 15 (20%) | 74 |  |  |
|  |  | C/C | 0 | 0 | 0 | 0 |  |  |
|  | rs117232607 | G/G | 535 (73%) | 152 (21%) | 37 (5%) | 724 | 1.96<br>(1.22-3.16) | 0.00475 |
|  |  | A/G | 43 (58%) | 16 (22%) | 15 (20%) | 74 |  |  |
|  |  | A/A | 0 | 0 | 0 | 0 |  |  |
|  | rs59666014 | C/C | 535 (73%) | 152 (21%) | 37 (5%) | 724 | 1.96<br>(1.22-3.16) | 0.00475 |
|  |  | A/C | 43 (58%) | 16 (22%) | 15 (20%) | 74 |  |  |
|  |  | A/A | 0 | 0 | 0 | 0 |  |  |

## Discussion

Our study demonstrates that targeted disruption of *Col4a1* specifically in adult brain microvessels is sufficient to induce CMBs that closely recapitulate human pathology. This novel approach offers unprecedented mechanistic insights into the role of BM integrity in cerebrovascular health and establishes a direct causal relationship between collagen IV deficiency and microbleed formation.

The strategy employed in this study offers distinct advantages over previous approaches to investigating Col4a1-related cerebrovascular pathology in adults. While earlier studies identified germline *Col4a1* mutations in mice causing porencephaly, ICH, and vascular lesions in multiple peripheral organs ^11,25^, these global mutations affect multiple tissues and developmental processes, complicating interpretation of the cerebrovascular phenotype, especially in adult subjects. Adult mice with germline *Col4a1* mutations that survived the perinatal period without obvious neurological abnormalities exhibited hemosiderin deposition, suggesting that CMBs can manifest as a consequence of these mutations ^11^. However, our targeted approach, which disrupts Col4a1 specifically in adult brain microvessels, clarifies the direct causal link between collagen IV deficiency and CMB formation, without the confounding effects of developmental or systemic factors. The temporal dimension of our intervention provides critical insights into BM function in mature vasculature. Previous work demonstrated that conditional *Col4a1* mutation induced after the developmental period (3 weeks postnatally) did not produce ICH or macroangiopathy, though CMBs were not specifically evaluated ^26^. In contrast, our targeted disruption of *Col4a1* at 8 weeks of age resulted in progressive accumulation of CMBs over several months, indicating that continuous BM integrity is essential for cerebrovascular health throughout adulthood and that its disruption leads to cumulative microvascular injury even in fully mature vessels.

Our ultrastructural analyses provide compelling mechanistic evidence linking collagen IV deficiency to compromised vascular structural integrity. EM revealed significant thinning of the brain vascular BM and reduced electron density in *Col4a1*-depleted microvessels compared to controls, with over 50% of vessels in the *Col4a1* group exhibiting BM thickness of 40 nm or less versus predominantly thicker structures (average ∼80 nm) in control animals. This structural deterioration establishes a direct mechanistic pathway whereby collagen IV depletion compromises the fundamental architecture of the vascular BM, leading to microvessel fragility and subsequent microbleed formation. The BM serves as a critical structural scaffold that maintains vessel wall integrity under physiological pressure ^27^, and its thinning may represent the primary mechanism underlying the microvascular vulnerability observed in our model.

The selective CMB phenotype in our model is particularly striking when compared to the complex presentation in human *COL4A1* mutations. Patients harboring *COL4A1* mutations exhibit diverse cerebrovascular disorders ranging from severe manifestations such as porencephaly and recurrent ICHs to subtle imaging abnormalities including CMBs, leukoaraiosis, and cerebral aneurysms ^10,11,14,28^. Recent clinical studies consistently report the presence of CMBs in the brain MRI of patients with *COL4A1* mutations, typically accompanied by other cerebrovascular features ^29,30^. These patients also frequently exhibit developmental brain abnormalities such as schizencephaly, focal epilepsy, and cortical malformations, along with systemic manifestations affecting the retina, kidney, heart, and muscles ^28,30^. The comparison of *Col4a1* mutation induction at various developmental stages in mice ^26^ suggests that disrupting *Col4a1* in adulthood likely does not compromise vascular integrity to a degree that would result in macroscopic ICH. Different features of CSVD pathology may have distinct mechanistic requirements. Specifically, collagen IV disruption in adult brain endothelial cells primarily compromises microvascular structural integrity, leading to microbleed formation without affecting vessel luminal patency or causing white matter ischemic lesions.

The existing CMB animal models include approaches induced by the accumulation of β-amyloid, cerebral hypoperfusion, genetic or dietary hypertension, and inflammation ^31,32^. Compared to these models, our model, created by targeted deletion of *Col4A1* specifically in brain microvessels, offers several distinct advantages. The model exhibits high penetrance across subjects and generates numerous CMBs within a practical timeframe (3-6 months), facilitating efficient quantitative evaluation. Crucially, it produces a pure CMB phenotype without confounding amyloid pathology, macrohemorrhages, or ischemic lesions that complicate interpretation in other models. The exceptional tunability of our system, with dose-dependent control over CMB number and size, would enable systematic investigation of how varying microbleed burdens influence behavioral and pathological outcomes. Furthermore, the progressive nature of CMB formation in our model aligns with longitudinal human clinical studies ^33,34^, providing a platform for evaluating therapeutic interventions aimed at preventing CMB progression.

The neurobehavioral deficits observed in our model, including cognitive impairment and motor incoordination, establish functional consequences of CMBs. These findings support clinical observations where increasing CMB burden correlates with cognitive decline ^33,34^. Notably, histological analyses revealed a distinctive pattern of glial activation, with widespread astrocytic reactivity extending well beyond microbleed sites while microglial activation remained localized. CLEM method demonstrated that microglial cells surrounded BECs and appeared to phagocytose electron-dense hemosiderin deposits within their plasma membranes. The difference in the spatial distribution between astrocytes and microglial cells suggests that astrocytic reactivity could be a more crucial mechanism connecting CMBs to neurological dysfunction, whereas microglial activation might indicate a more localized response to iron deposition. Reactive astrocytes can release various cytokines and alter their support functions for neurons and synapses ^35,36^, potentially disrupting network activity. This widespread astrocytic response could explain how multiple small microbleeds can collectively impact cognitive and motor functions despite their limited individual spatial extent. In *Col4a1*-deleted animals, increased ferritin immunoreactivity suggested a higher iron load in brain tissue surrounding CMBs, which may be associated with widespread activation of astrocytes.

The analysis of human genetic data showed a significant correlation of CMB number with genetic variants of *Col4a1* and *TIMP2*, underscoring the clinical relevance of our CMB model generated by targeted *Col4a1* deletion. Collagen IV in the vascular BM is degraded by various MMPs, particularly by MMP2 and MMP9 ^37^. Our data implicate specific involvement of the collagen IV-MMP2 pathway in the pathogenesis of sporadic CMB formation. TIMP2 is a key endogenous inhibitor of matrix MMPs, particularly MMP2 ^24^, and by inhibiting MMP2, it helps prevent excessive degradation of collagen IV and other basal lamina components, maintaining the integrity of brain microvessels ^38^. The identification of 4 polymorphisms in *TIMP2* strongly associated with CMB burden suggests that dysregulation of the homeostatic balance between BM synthesis and degradation may be a common mechanism underlying CMB formation in humans. A recent study showed that MMPs were upregulated in *COL4A1* mutant patient-derived cells, which correlated with ECM deposition and increased cell death ^39^, supporting the implication of homeostatic imbalance in Collagen IV-related vasculopathy.

In conclusion, our CMB model generated by targeted *Col4a1* deletion in brain microvessels offers novel insights into the specific contribution of BM disruption to CMB formation and associated neurological impairments. The convergence of experimental findings and human genetic evidence suggests that similar mechanisms underlie sporadic CMBs in aging populations. Thus, our model may represent an invaluable tool for elucidating precise pathogenic mechanisms and developing therapeutic strategies to halt the progression of CMB formation. This model may also have important implications for understanding amyloid-related imaging abnormalities-hemosiderin (ARIA-H) observed in patients receiving amyloid-targeting therapies for Alzheimer’s disease ^40^. Our findings suggest that underlying BM vulnerability, potentially influenced by genetic factors such as TIMP2 variants or other alterations in the mechanisms regulating its degradation ^41^, could predispose certain patients to develop ARIA-H.

## MATERIALS AND METHODS

### Animals

CRISPR/Cas9 knock-in mice (B6(C)-*Gt(ROSA)26Sor^em^*^1^.^1^(CAG–cas9*,–EGFP)*^Fezh^*/J) were purchased from Jackson Laboratory (Strain #:026179). Genotyping was carried out according to the vendor’s instructions. All animal experiments were approved by the Ajou University Medical Center Institutional Animal Care and Use Committee (IACUC 2024-0028). All experiments involving animals were conducted in accordance with the ARRIVE guidelines. All mice were maintained on a regular diurnal lighting cycle (12:12 light:dark) with ad libitum access to food and water. The maximum caging density was five mice from the same litter and sex starting from weaning.

### *In vitro* screening of mouse col4a1 targeting gRNA

To knockout mouse *Col4a1*, we designed SpCas9 (Cas9 derived from *Streptococcus Pyogene*) guide RNAs that can target the early coding region of mouse *Col4a1* with minimal off target potential (gRNAs with 0 off-targets in the reference genome up to 2 base pair mismatches were selected). From these criteria, we designed total 9 gRNAs in either exon 1 or exon 21. sgRNA targeting the mouse *Rosa26* locus was utilized as a control. Individual sgRNA sequences were cloned into the pRGEN-Cas9-CMV/T7-RFP plasmids. These were then electroporated into NIH3T3 mouse fibroblast cell lines (ATCC, #CRL-1658) using a Neon Electroporation System (Thermo-Fisher). Secondary screening was conducted using lentivirus-mediated gene editing techniques. For this, the candidate sgRNAs were cloned into a lentiviral vector (pLV-mCherry-U6 backbone; Vectorbuilder), and lentiviral particles were treated to cultured primary BECs obtained from the Cas9 knock-in mice. Based on secondary screening, a lead sgRNA sequence was chosen (Fig. 1A) and cloned into AAV-U6-CMV-mCherry or AAV-nano luciferase-U6 plasmid.

### Targeted deep sequencing

We performed targeted deep sequencing to quantitatively measure gene editing efficiency as previously described ^42,43^. Briefly, the on-target region was PCR-amplified from gDNA extracted from transfected cells using Phusion polymerase (New England BioLabs). The resulting PCR amplicons were then subjected to paired-end deep sequencing using MiSeq (Illumina). Data from deep sequencing were analyzed using the online Cas-Analyzer tool (www.rgenome.net) ^44^. Indels located 3 base pairs upstream from the protospacer-adjacent motif (PAM) sequence were considered mutations caused by Cas9.

### Preparation and injection of AAV-BR1 particles

The AAV BR1 serotype viral particles containing GFP expression cassette (AAV-BR1-GFP) were purchased from SignaGen Laboratories (Frederick, MD). AAV-BR1 viral particles containing sgRNA sequences were produced by PackGene (Houston, TX). A 100 μl volume of PBS containing different concentrations of AAV-BR1 (1.0, 2.5, and 5.0 × 10^13^ GC/kg of body weight) was injected into the retrobulbar venous sinus using the previously described method ^45^. The mice were anesthetized with 2.5% isoflurane, and then a 31G insulin syringe loaded with the AAV solution was positioned at a 45° angle to the sagittal plane and parallel to the horizontal plane of the mouse’s anatomical axis to locate the retro-orbital venous sinus. The syringe needle was gently advanced until it touched the orbital bone structure. The needle was then slightly retracted, and the plunger was carefully depressed for approximately 2 seconds to deliver the solution. The syringe was slowly removed 30 seconds after the injection. Animals were closely monitored to ensure there was no bleeding or bulging of the eyes during the infusion.

### Cell culture

Primary BECs were isolated from 8-week-old C57BL/6 or Cas9 transgenic mice following the previous study with slight modifications ^46^. The mice were anesthetized, and their brains were gently dissected and collected in ice-cold PBS. Care was taken to remove meninges and any visible white matter. The cerebral cortices were cut into small pieces using sterile razor blades. The cortical tissues were then homogenized with a tissue grinder and centrifuged at 1,350g for 5 minutes. The resulting pellet was resuspended in a dextran solution, centrifuged at 6,100g for 10 minutes, and digested in DMEM containing collagenase/dispase, TLCK (Tosyl-L-lysyl-chloromethane hydrochloride, trypsin inhibitor), and DNase I at 37°C for 75 minutes until being terminated by centrifugation at 1,350*g* for 5 min. The pellet was resuspended in DPBS, centrifuged at 1,350g for 5 minutes, and then resuspended in full medium (DMEM-F12 containing 10% FBS, ECGS, heparin, 1% antibiotic/antimycotic, and L-glutamine). The resuspended microvessel fragments were seeded on collagen type IV-coated 35 mm plastic dishes. The primary BECs originating from microvessel fragments were maintained with 5% CO2/95% air at 37°C in full medium containing puromycin (8 µg/ml) for the first 2 days to obtain a pure culture of BECs. Unlike pericytes, BECs can survive due to their high expression of efflux pumps that scavenge the intracellular toxicity generated by puromycin [3]. The BECs were changed with full medium 2 times per week, split after 7-10 days, and cultured for 3 days before being used in the experiments.

### Immunocytochemistry

The cells were cultured on coverslips, fixed with 4% paraformaldehyde in phosphate buffered saline (PBS) at 25°C for 10 minutes, and permeabilized by incubating in PBS containing 0.1% Triton-X100 for 10 minutes before blocking for 1 hour with PBS containing 1% BSA and 0.1% Triton-X100. All primary antibodies were incubated overnight for staining at 4°C in the same blocking solution. Immunoreactive proteins were visualized by incubation with Alexa Fluor 488- or Alexa Fluor 546-conjugated secondary antibodies (Invitrogen). Cells were mounted with a DAPI-containing mounting solution and observed under a confocal microscope (Carl Zeiss, Jena, Germany).

### Quantitative real time RT-PCR

Total RNA was isolated using RNAiso Plus (TaKaRa, Otsu, Japan), and cDNA was synthesized using avian myeloblastosis virus reverse transcriptase (New England Biolabs) and oligo(dT) primers (Promega, Madison, WI, USA), according to the manufacturer’s instructions. For quantitative real-time PCR, we used a Thermal Cycler Dice Real Time System (TaKaRa) with SYBR Premix Ex Taq master mix (TaKaRa) in accordance with the manufacturer’s instructions. The *Col4a1* and *Gapdh* primers for qPCR (Bioneer, Daejeon, Korea) are as follows: mouse *Col4a1* primer, 5’-ATG GCT TGC CTG GAG AGA TAG G -3’ (forward) and 5’-TGG TTG CCC TTT GAG TCC TGG A -3’ (reverse); mouse *Gapdh* primer, 5’-CAT CAC TGC CAC CCA GAA GAC TG -3’ (forward) and 5’-ATG CCA GTG AGC TTC CCG TTC AG -3’ (reverse).

### Western blotting

The BECs were lysed in RIPA buffer containing 50 mM Tris-HCl (pH 7.4), 150 mM NaCl, 0.5% NP-40, 1 mM DTT, 1 mM PMSF, a protease inhibitor cocktail, a phosphatase inhibitor cocktail, 5 mM NaF, 1 mM sodium vanadate, and 1 mM EDTA. The protein concentration in lysates was determined using the Bio-Rad protein assay according to the manufacturer’s instructions. Subsequently, 10 μg of proteins were separated by sodium dodecyl sulfate-polyacrylamide gel electrophoresis (SDS-PAGE) on 10% polyacrylamide gels and transferred to a nitrocellulose membrane. The membranes were probed using the specified primary and secondary antibodies and developed using an enhanced chemiluminescence detection kit (WESTSAVE Gold; AbFrontier, Seoul, Korea).

### Magnetic resonance imaging of the mouse brain

The animals were given 1-1.5% isoflurane anesthesia during the MRI procedure. Their respiration rate (80-120 breaths per minute) and body temperature (36.5 ± 0.5°C) were monitored throughout the experiments. The MRI scans were carried out using a horizontal bore 9.4T/30 cm Bruker BioSpec MR system (Billerica, MA, USA) at the Center for Neuroscience Imaging Research (CNIR) of Sung Kyun Kwan University (SKKU). An actively shielded 12 cm diameter insert with a maximum strength of 66 gauss/cm and a rise time of 141 μs was used. For excitation, a quadrature birdcage coil (86 mm inner diameter) was used, while detection utilized an actively decoupled planar surface coil (10 mm inner diameter) positioned on top of the mouse head. A T2*-weighted MRI was obtained using the FLASH (Fast low angle shot) sequence with the following parameters: repetition time (TR)/echo time (TE) = 600/8 ms, number of excitations (NEX) = 6, field of view = 14 (readout) x 11 (phase encoding) mm^2^, flip angle = 60°, matrix = 350 x 275, in-plane resolution = 40 x 40 μm2, slice thickness = 500 μm, with 25 contiguous slices without gap in the coronal plane. We selected four coronal sections based on distances from the bregma to count the number of cerebral microbleeds on MRI in specific anatomical locations within the brain. These locations were: +1.98 mm for the prefrontal cortex, +0.5 mm for the sensorimotor cortex and striatum, -2.06 mm for the hippocampus and thalamus/hypothalamus, and -4.04 mm for the brainstem. We counted all hypointense foci suggestive of paramagnetic signals at these specific anatomical locations, excluding any spots with a diameter less than 0.01 mm. Additionally, hypointensity signals located on the meninges and ventricular spaces were not included in the count.

### Behavioral assessments

The mice underwent a series of behavioral tests in a quiet room with infrared lighting. Mice were randomly divided into either AAV-BR1-sgRNA-*Col4a1* or AAV-BR1-sgRNA-*Rosa26* injection group. After the injection, they were assigned new identification codes to ensure blind evaluation during the course of behavioral tests. These tests included novel object recognition (NOR), Y-maze, and rotarod. A video tracking system (SMART 3.0, Panlab) was utilized to automatically measure motor behavior during testing. The NOR test was conducted based on previous reports with a slight modification ^47,48^. One day prior to the actual testing, all mice were exposed to an empty chamber measuring 400 × 400 mm for 20 minutes. During this habituation session, we measured the total distance of locomotion as an index of the overall locomotor activity (open field test). During the NOR test, there were two sessions: “familiarization” and “test,” each lasting 10 minutes. In the familiarization session, two copies of a familiar object were placed five centimeters away from the walls in opposing corners of the chamber. During the test session, a duplicate of the familiar object and a new object were placed in opposing corners of the chamber. Between the two sessions, the apparatus and all objects were cleaned with a 70% ethanol solution to eliminate any odor cues. Object exploration is defined as the behavior of sniffing or touching the object while observing it, with the distance between the nose and the object being less than 5 cm. The discrimination index for each animal was calculated using the formula: Discrimination index (%) = (time spent exploring the new object / time spent exploring both objects) × 100. For the Y-maze test, we modified the protocol of our previous study using rats to adapt it for mice ^49^.

The apparatus consisted of three identical arms made of white plastic, each measuring 325 mm in length, 70 mm in width, and 150 mm in height. The mice were positioned at the center of the maze and given 10 minutes to freely explore the three arms. The entries into each arm were manually determined based on the video recording. An alternation triplet was considered if there was a consecutive sequence of entries into all three arms. For example, if the sequence of entries into arms labeled A, B, and C is ABCBCABC, there are 4 alternation triplets (ABC, BCA, CAB, and ABC). The percent alternation triplet was calculated using the formula: Alternation triplet (%) = {number of alternation triplets / (total number of arm entries – 2)} × 100. The rotarod apparatus for mice (Ugo Basile, Comerio, Italy) was used to assess the coordination and balance of the mice. Before the actual testing, the mice were familiarized with the rotarod apparatus over 3 days. During this habituation phase, each mouse was placed on the rotarod at a constant speed of 4 rpm for up to 3 minutes, and this was repeated 4 times each day. The testing phase lasted for 3 consecutive days, with 3 trials each day. During each trial, the rotarod speed was gradually increased from 4 to 40 rpm over 5 minutes. The time taken to fall from the apparatus was recorded for each trial, and the average time from the three trials of each day was calculated. To minimize differences between subjects, the average time from the 3-day trials was normalized by the average time before the AAV injection.

### Intravital imaging using a two-photon microscope

The mice were anesthetized by intraperitoneal injection of a ketamine (90mg/kg) and xylazine mixture (10 mg/kg) and the skin above the skull was removed. A circular area of the skull with a 4 mm diameter (positioned at 1.5 mm posterior and 1.3 mm lateral to the bregma was removed to expose the dura mater. A coverslip (5 mm) was placed over the opening and secured in place with cyanoacrylate glue, and dental acrylic cement was used to cover the skull surface. Warm saline (37 °C) was continuously supplied during imaging to prevent bleeding and drying of the exposed brain surface. Imaging was performed using an upright LSM980 microscope (Carl Zeis) equipped with a MaiTai DeepSee femtosecond-pulsed laser (Spectra-Physics, Santa Clara, CA, USA) with an excitation wavelength of 800 nm. The brain vessels were labeled by injecting 100 µL of 150 kDa Fluorescein isothiocyanate-dextran (FITC-dextran, 30mg/ml, Sigma Aldrich) via the retro-orbital route. The 2-photon microscopic images were acquired as z-stacks with 0.5 μm step size between each focus plane. The z-projections and 3D image reconstruction were performed using IMARIS software (ver. 9.8).

### Histological processes and immunohistochemistry

The mice were transcardially perfused with PBS (pH 7.4) and 4% paraformaldehyde in 0.1 M phosphate buffer (4% PFA/PB). The whole brain was immersed in 4% PFA/PB for 2h at 4°C and cut into 30 μm coronal sections using a vibratome (VT1200 S, Leica Microsystems, Wetzlar, Germany). The brain sections were evaluated histologically using hematoxylin and eosin (H&E) staining and non-heme iron histochemistry (Prussian blue reaction). Coronal sections were placed on a slide, dehydrated, rehydrated, and then stained with hematoxylin (Sigma). The slides were rinsed in tap water, counterstained with eosin, dehydrated, and mounted using a xylene-based Permount solution. To conduct Prussian blue staining, 30-μm sections were immersed in a solution containing 5% potassium ferrocyanide in aqueous hydrochloric acid for 5 minutes. Following incubation, the sections were rinsed in water and subsequently counterstained with nuclear fast red (Sigma-Aldrich, Burlington, MA). The number of Prussian blue stained deposits with an obvious purple-blue color was counted within specific anatomical locations using a 10× objective lens. Very small spots with a diameter smaller than 10 μm, occasionally observed within intracellular spaces, were excluded from the counting. Free-floating coronal brain sections were permeabilized with PBS containing 0.3% Triton X-100 (0.3 % PBST) and blocked with 0.3 % PBST containing 10% normal goat serum. Then, they incubated at 4°C overnight with the following primary antibodies: rat monoclonal anti-CD31 (1:50, #550274, BD Pharmingen), rabbit polyclonal anti-Iba1 (1:500, #019-19741, Fujifilm Wako), rabbit polyclonal anti-CD45 (1:500, #NB100-77417, Novus Biologicals), mouse anti-GFAP (1:250, #sc-33673, Santa Cruz Biotechnology), and rabbit polyclonal anti-COL4A1 (1:50, #ab6548, Abcam). After rinsing 3 times with 0.3 % PBST at RT, the sections were incubated with the appropriate secondary antibodies labeled with fluorescence for 1h and mounted on glass slides for confocal microscopic observation (LSM900, Carl Zeiss). The expression of the ferritin (light and heavy chain) in the brain sections was determined using rabbit polyclonal anti-ferritin light chain (1:250, #ab69090, Abcam) and rabbit monoclonal anti-ferritin heavy chain antibody (1:100, #ab183781, Abcam). The reactions were performed using indicated primary antibodies overnight at room temperature, washed, and incubated with secondary biotinylated antibody and streptavidin-peroxidase complex for 1hr at RT. NovaRed (Vector Laboratories, Newark, CA) was used as a chromogen for 5 min at RT and counterstained with Meyer’s hematoxylin. The negative control was performed by omitting the primary antibody. Images were analyzed with Axioscan Z1 Microscope Slide Scanner (Carl Zeiss). To image hemosiderin-laden IBA1+ microglia, CD45+ macrophages, and GFAP+ astrocytes, the sections were first incubated with anti-Iba1, anti-CD45, and anti-GFAP antibodies. After washing, the sections were then incubated with secondary biotinylated antibodies and streptavidin-peroxidase complex for 1h at RT. Then, the sections were transferred to slide glasses, incubated with NovaRed, and stained using the Prussian blue reaction method.

### Electron microscopic analysis

For correlative light-electron microscopy (CLEM), harvested brain tissues were initially trimmed into 1 mm^3^ blocks and post-fixed with 4% paraformaldehyde (PFA) for 3 hours. Subsequently, tissues were immersed in 30% sucrose solution in 0.1M phosphate buffer (PB) at 4°C until fully sunk, after which they were transferred into a 2.3 M sucrose solution in 0.1 M PB and incubated for 24 hours at 4°C. Following cryoprotection, samples were preserved in liquid nitrogen as previously described ^50^. Semi-thin cryosections (2 μm thick) were then prepared at -100°C using a glass knife in a Leica EM UC7 ultramicrotome equipped with an FC7 cryochamber (Leica). The sections were immunolabeled with mouse monoclonal anti-mCherry antibody (1:200, Sigma-Aldrich, # SAB2702295), goat anti-human/mouse/rat CD31/PECAM-1 polyclonal antibody (1:250, R&D Systems, # AF3628), rabbit polyclonal anti-Iba1 antibody (1:300, FUJIFILM Wako Pure Chemical Corporation, # 019-19741) and chicken polyclonal anti-GFAP antibody (1:700, Millipore, # AB5541). Antibody staining was visualized using Cy3-conjugated donkey anti-mouse IgG (1:2000, Jackson ImmunoResearch, # 715-166-150), Alexa Fluor® 488-conjugated donkey anti-goat IgG (1:300, Invitrogen, # A11055, Thermo Fisher Scientific), Alexa Fluor® 647-conjugated donkey anti-rabbit IgG (1:300, Invitrogen, # A31573, Thermo Fisher Scientific) and Cy5-conjugated donkey anti-chicken IgG (1:1000, Jackson ImmunoResearch, # 703-175-155). Sections were labeled with DAPI for 10 minutes. Immunolabeled sections were photographed using a confocal microscope (LSM900 with Airyscan), post-fixed with 2.5% glutaraldehyde in 0.1 M phosphate buffer followed by 1% osmium tetroxide (OsO_4_), and embedded in Spurr’s resin (Electron Microscopy Sciences, #14300). The processed samples were subsequently analyzed using scanning electron microscopy (Regulus 8220, Hitachi). For conventional transmission electron microscopy, mouse brain tissues were fixed using 2% paraformaldehyde and 2% glutaraldehyde in 0.1 M sodium cacodylate buffer. The samples were stained with 1% OsO4, dehydrated in a series of graded ethanol, and then embedded in resin EMbed 812 (EMS) for 48 hours at 60°C. Ultrathin sections were cut using an EM UC7 ultramicrotome (Leica). The sections were stained with lead citrate and uranyl acetate and analyzed under a Zeiss Sigma 500 electron microscope at the Three-Dimensional Immune System Imaging Core Facility of Ajou University. Quantification of the BM thickness was conducted following a previously reported protocol with slight modifications ^51^. We selected microvessels with rounded lumens that displayed clear BM structures. For each capillary, we randomly selected three distinct regions of the BM for analysis of thickness. We then measured the distances between the inner and outer edges of these regions using the perpendicular measurement function in ImageJ software.

### Evaluation of CMBs in human MR images

The study using human data was approved by the Ajou University Institutional Review Board (IRB) (AJOUIRB-EX-2024-156). All human data, including MR images, clinical information, and genomics data using SNP microarray chips were obtained from the Korean government initiative designated as the Biobank Innovations for Chronic Cerebrovascular Disease with ALZheimer’s Disease Study (BICWALZS), a multi-center collaborative project led by Ajou University Hospital. The participants in this project were individuals (Korean, east Asian ethnicity) who visited specialized neurology or psychiatric outpatient memory clinics for the evaluation of subjective memory impairment. Details on the baseline clinical and biomarker characteristics of the BICWALZS were previously reported ^21^. MRI scan data were obtained using a 3.0T MR scanner. CMBs were assessed with either the gradient echo T2* or susceptibility-weighted imaging (SWI) sequence images. MR images from 849 out of a total of 1308 participants were deemed to have suitable quality for CMB counting. 13 study participants revealed multiple CMBs, ranging from 15 to 154, with a median of 46, predominantly located in the cerebral cortex, suggesting the presence of amyloid angiopathy ^52^. Because amyloid angiopathy is strongly associated with genetic variants related to Alzheimer’s disease, we excluded the data from these 13 subjects. CMBs were defined as well-delineated, round, or ovoid hypodense foci measuring less than 10 mm in diameter with blooming artifacts ^17^. Hypointensity structures within the subarachnoid space or areas of symmetric hypointensities within the globus pallidus were not counted as CMBs because they are highly likely to represent adjacent pial blood vessels and calcifications, respectively ^53^. We also followed the recommended criteria for CMB identification and took special care not to include MRI signals that could confound and mimic CMBs ^18^ All the MR images were independently reviewed by three researchers (H.K., H.K.G., and S.S.S.) under the supervision of two board-certified neurologists (S.Y.G. and B.G.K.). The images where the evaluation between the three researchers was not consistent were finally reviewed by the two neurologists until agreement was reached.

### SNP genotyping and association analysis with CMB phenotype

SNP genotyping was conducted using the Korea Biobank Array (Affymetrix Axiom KORV1.1-96 Array, Thermo Fisher Scientific) at DNA Link Inc. (Seoul, South Korea). DNA samples that did not meet the specific criteria ^54^ were excluded from further analyses. As a result, SNP genotyping data could not be obtained from 25 out of 836 subjects whose MRI data were available for CMB analysis, and the final number of subjects included in the study on the relationship between SNP and CMBs on MRI was 811. The quality control processes for SNP genotyping were previously described in detail ^54^. Briefly, SNPolisher was utilized to exclude low-quality markers based on poly high resolution, mono high resolution, and no minor homozygosity. Study-specific association analyses for sequencing the data of the 811 subjects were performed using linear regression for the number of microbleeds, assuming an additive genetic model. We systematically excluded variants with minor allele frequency (MAF)[>[5% and a *p* value < 1.0 × 10^-5^ for the Hardy-Weinberg equilibrium test, which has been the conventional threshold for a common variant, using dbSNP database (version 150). We screened the relationship between CMBs and SNPs using PLINK (version 1.9), an online tool for genome-wide association studies and genetic association management through multi-dimensional scaling (MDS) analysis ^55^. The covariates were age, sex, body mass index (BMI), diabetes mellitus, dyslipidemia, hypertension, and history of previous stroke. The protein-protein interaction network of 100 curated genes, of which SNPs were most robustly correlated with CMB number, was created using STRING database (https://string-db.org/, version 12.0). Individual SNPs that showed a significant correlation with CMB number were further analyzed by calculating a contingency table based on the genotype data and trichotomized categories of CMB numbers. A chi-square test was used to assess the statistical significance of the association between genotype and the proportion of study subjects in different CMB number groups.

### Statistical analysis

Statistical analyses were performed using SPSS 25.0 and GraphPad Prism 10.2 software. Student’s t-test (two-tailed) was used to compare mean values between two independent groups, while one-way ANOVA was employed for three or more independent groups, followed by Tukey’s *post hoc* analysis. For monthly comparisons of cognitive and motor performance over six months, repeated measures two-way ANOVA was conducted, followed by Bonferroni *post hoc* analysis to assess group differences at each time point. The two-sample Kolmogorov-Smirnov test was used to evaluate statistical differences in the distribution of CMB diameter and BM thickness observed in EM images. Details on SNP statistical analysis are provided in the corresponding section.

## Data availability

All data supporting the findings of this study are available within the paper and its Supplementary Information. The raw results of the GWAS analysis are available upon request. The complete dataset, which includes MRI analysis and genotyping results, is also available upon request. However, the Ajou University IRB will review all requests related to human data to determine whether each request is subject to confidentiality restrictions.

## Acknowledgments

We thank the Three-Dimensional Immune System Imaging Core Facility of Ajou University for providing intravital imaging services and the Institute for Basic Science (IBS) Center for Neuroscience Imaging Research (IBS-R015-D1) for its support, including MRI time and professional technical assistance.

## Author contributions

HK, JYL, and BGK planned research and designed experiments. HK performed and led the majority of experiments. YS, HK, SSS, JL, and HL participated in the acquisition of cellular, molecular, and behavioral data from animal studies. JWH and TRR contributed to the generation of electron microscopy results. HK, SSS, SK, and BGK reviewed and analyzed human MRI data. JYC, HWR, and SJS participated in the design of human data collection and database management. HK, GTK, SKC, HSJ, SYJ, and BGK designed the SNP experiment and conducted statistical analysis. HK, KIL, JYL, and BGK interpreted all the data and contributed to drawing the main conclusions. HK, JYL, HSJ, and BGK wrote the manuscript. JYL and BGK approved the manuscript submission.

## Funding information

This research was supported by the National Research Foundation of Korea (NRF) research programs (RS-2019-NR040055, RS-2021-NR056919, and RS-2023-00244748 to BGK, and RS-2023-00245169 to HK), by Korea Initiative for fostering University of Research and Innovation (KIURI) Program of the National Research Foundation (NRF) funded by the Korean government (NRF2021M3H1A104892211), and by 2023 intramural research fund of Ajou University Medical Center (M2023C046000102 to HK).

## Competing Interests statement

All participating authors have no competing interests as defined by Nature Portfolio, or other interests that might be perceived to influence the results and/or discussion reported in this paper.

## Figure legends

**Figure S1*. In vitro* screening of sgRNA targeting the mouse *Col4a1* gene**

**(A, B)** Targeted deep sequencing and **(C)** quantitative RT-PCR analysis of NIH-3T3 cells edited by candidate *Col4a1*-targeting sgRNAs. **(D, E)** Quantitative RT-PCR **(D)** and Western blot analysis **(E)** of primary brain endothelial cells (BECs) obtained from Cas9 overexpressing transgenic mice. BECs were transfected with sgRNAs #4 and #5, targeting exon 1 of *Col4a1*, using the electroporation system. *R*: gRNAs sgRNAs targeting the mouse *Rosa26* locus. Beta actin was used as an internal loading control.

**Figure S2. Specificity of AAV-BR1 transduction.**

**(A)** Representative immunohistochemical (IHC) images of brain slices obtained from 8-week-old mice injected with AAV-BR1-GFP. IHC staining was performed using antibodies against CD31 (brain EC marker), GFAP (astrocyte marker), IBA-1 (microglial marker), and MAP2 (neuronal marker). Scale bars = 50 µm. **(B)** Representative IHC images of liver, heart, and lung slices obtained from 8-week-old mice injected with AAV-BR1-GFP. Scale bars = 50 µm.

**Figure S3. No evidence of ischemic injury in animals with targeted *Col4a1* deletion.**

**(A)** Representative images of the coronal brain sections stained with Eriochrome cyanine RC. Rectangular regions containing the corpus callosum and the cingulum are magnified on the right side. **(B)** Representative images of the coronal brain sections immunostained with antibodies against myelin basic. Rectangular regions containing the corpus callosum and the cingulum are magnified on the right side. Scale bars = 100 µm.

**Figure S4. Iron deposition does not influence the integrity of neurons and myelinated fibers.**

**(A, B)** Representative images of the coronal brain sections that were subjected to immunohitochemical staining with ether NeuN **(A)** or myelin basic protein (MBP) **(B)**. The sections were then processed with Prussian blue staining to visualize hemosiderin deposits. Rectangular regions are magnified in the lower panels. All scale bars = 20 µm.

**Figure S5. Quantification of general motor activity and body weight.**

**(A-C**) Quantification of general motor activity in an open field. The total distance traveled by animals during the test session was quantified as a measure of general motor activity. **(D-F)** Quantification of body weight. Animals were injected at the age of 8 weeks. The test was conducted before the injection and then once a month for six months following the injection. N = 14 (7 per gender) and 18 (9 per gender) for control sgRNA against Rosa26 and targeting Col4a1, respectively.

**Figure S6. Human MR images representing three groups with different CMB burdens.**

Representative MR images from the study subjects that were classified into three groups depending on CMB burden: no CMBs (Group I), 1-4 CMBs (Group II), and ≥ 5 CMBs (Group III). MR images were acquired using either T2*-weighted gradient echo (GRE) or susceptibility-weighted imaging (SWI) sequences. Arrowheads indicate isolated CMBs in Group II.

**Figure S7. A Manhattan plot of the genome-wide association study of CMB burdens.**

**Figure S8. Regional association plots of *TIMP2* variants significantly associated with CMB burden.**

**(A)** Regional association plot with rs3744787 as the reference SNP (purple diamond) spanning a 50 kb genomic window. Association summary statistics are presented as - log[[(P-values) versus chromosomal position. **(B)** Fine-mapping view centered on rs118160201 as the reference variant (purple diamond). Three linked SNPs (rs117232607, rs118160201, and rs59666014) are clustered within a 1-kb region. Data points are colored according to their linkage disequilibrium (r²) with the reference SNP using 1000 Genomes Project Asian ancestry data. Red circles indicate strong correlation (r² > 0.8) with the reference variant for rs117232607 and rs59666014. Black dashed lines in **(A)** delineate the genomic boundaries magnified in **(B)**. Regional plots generated using LocusZoom (locuszoom.org).

**Movie S1. A time-lapse video clip visualizing the leakage of fluorescently labeled dextran from brain vessels in an animal injected with AAV-BR1-sgRNA-*Col4a1*.**

**Movie S2. Video clips showing 3D reconstruction of high-resolution stacks of the cerebral blood vessels visualized by intravenous injection of FITC-labeled dextran.**

## Notes

### Competing Interest Statement

The authors have declared no competing interest.

